# The genomic basis of domestic colonisation and dispersal in Chagas disease vectors

**DOI:** 10.1101/2021.04.27.441467

**Authors:** Luis E Hernandez-Castro, Anita G Villacís, Arne Jacobs, Bachar Cheaib, Casey C Day, Sofía Ocaña-Mayorga, Cesar A Yumiseva, Antonella Bacigalupo, Björn Andersson, Louise Matthews, Erin L Landguth, Jaime A Costales, Martin S Llewellyn, Mario J Grijalva

## Abstract

The biology of vector adaptation to the human habitat remains poorly understood for many arthropod-borne diseases but underpins effective and sustainable disease control. We adopted a landscape genomics approach to investigate gene flow, signatures of local adaptation, and drivers of population structure among multiple linked wild and domestic population pairs in *Rhodnius ecuadoriensis*, an important vector of Chagas Disease. Evidence of high triatomine gene flow (F_ST_) between wild and domestic ecotopes at sites throughout the study area indicate insecticide-based control will be hindered by constant re-infestation of houses. Genome scans revealed genetic loci with strong signal of local adaptation to the domestic setting, which we mapped to annotated regions in the *Rhodnius prolixus* genome. Our landscape genomic mixed effects models showed *Rhodnius ecuadoriensis* population structure and connectivity is driven by landscape elevation at a regional scale. Our ecologically- and spatially-explicit vector dispersal model enables targeted vector control and recommends spatially discrete, periodic interventions to local authorities as more efficacious than current, haphazard approaches. In tandem, evidence for parallel genomic adaptation to colonisation of the domestic environment at multiple sites sheds new light on the evolutionary basis of adaptation to the human host in arthropod vectors.

## Main

The process by which insect vectors of human diseases adapt to survive and breed in human habitats is fundamental to the emergence and spread of vector-bone disease (e.g., *Aedes aegypti*^1^). Relatively modest changes in vector host preference between ancestral (wild) and derived (domesticated) forms can drive devastating epidemics that result in millions of deaths^2^. Understanding the evolution and genetic bases of traits associated with domestication in disease vectors is, therefore, paramount and could inform control efforts and reveal the epidemic potential for new vector species^3,4^. Furthermore, an accurate definition of landscape functional connectivity (the level at which the landscape heterogeneity facilitates or impedes an organism’s movement from, and to, different habitat patches^5^) can shed light on the drivers of vector dispersal, and even assist in identifying poorly connected or isolated areas that can be easily targeted by eradication interventions^6–8^.

Triatominae (Hemiptera: Reduviidae) are a group of hematophagous arthropods that transmit *Trypanosoma cruzi*, the parasite that causes Chagas disease, a fatal parasitic infection afflicting > 7 million people in Latin America^9^. Eradication of ‘domesticated’ triatomines has been the mainstay of disease control in the past (e.g., *Triatoma infestans*^10^, *Rhodnius prolixus* and *Triatoma dimidiata*^11^). However, wild (e.g., *T. infestans*^12^ and *R. prolixus*^13^) and/or secondary competent species of triatomines (e.g., *Triatoma sordida*^14^, *Triatoma maculata* and *Rhodnius pallescens*^15^, *Panstrongylus howardi*^16^ and *P. chinai*^17^) can occupy empty domestic niches and continue to jeopardise Chagas disease control strategies.

Colonisation of the domestic niche may involve multiple, independent evolutionary processes across the geographic distribution of a given vector species^18,19^, analogous to parallel trophic speciation observed in other arthropods^20^. Alternately, domestication of zoonotic parasites and their vectors may result from a single or limited number of independent colonisation events, followed by rapid and widespread dispersal within the domestic setting^21,22^. Domestication of a given species may also represent a combination of these two scenarios, where multiple domesticated lineages serially introgress with wild lineages over evolutionary time, as has been elegantly demonstrated through analysis of the genomes of the domestic pig^23^. Disentangling these different scenarios in triatomine species, and their important implications for disease control, has been challenging due to a lack of genomic resources for these organisms which are only recently becoming available^24–26^. With adequate genomic tools; however, the occurrence of domestic colonisation can be established, and its underlying mechanisms unveiled. Parallel colonisation events explored using models of ‘adaptation with gene flow’ (e.g., ^27^) can exploit standard population genetic metrics and theory to make generalisations about the genomic basis of adaptations (e.g., ^20^) and reveal fundamental traits associated with the domestic niche.

*Rhodnius ecuadoriensis* is the major vector for Chagas disease in Ecuador and Northern Peru^28^. Both domestic and wild populations of this species exist throughout its range^29^. Preliminary morphological and genetic evidence suggests some gene flow of *R. ecuadoriensis* between domestic and wild ecotopes^30,31^. By comparison, genetic studies of *T. cruzi* infecting the same vectors in Ecuador have shown strong to moderate differentiation between wild and domestic isolates^32,33^. As such there is a lack of a clear understanding of the micro and macro-evolutionary and ecological forces shaping vector domestic adaptation and dispersal capabilities, and those of the parasites they transmit. Morphometric studies have attempted to develop phenotypic markers in triatomines associated with domestic or wild ecotopes with little (e.g. ^34^) to moderate (e.g., ^35^) success. Therefore, domestication in triatomines has become a rather qualitative concept^36^ with urgent need for quantitative foundations.

Our study represents a first attempt to accurately quantify genomic signatures of domestication of triatomine species, as well as landscape drivers of vector dispersal. We use a reduced-representation sequencing approach (2b-RADseq) to recover genome-wide SNP variation in 272 *Rhodnius ecuadoriensis* individuals collected across ecological gradients in Loja, Ecuador. We find strong evidence of gene flow between domestic and wild ecotopes and signatures of local adaptation in some genomic regions. Furthermore, we provide substantial evidence that triatomine dispersal is fundamentally restricted by landscape elevation. Our findings suggest frequent and spatially targeted interventions, to cope with high gene flow and fragmented populations, are necessary to suppress Chagas Disease transmission in Loja. Moreover, discovery of signatures of local adaptation shed the first light on the genomics basis of domestication in triatomines.

## Results

### Recovery of SNP markers from 272 *Rhodnius ecuadoriensis* SNP specimens

Our CspCI-based 2b-RAD protocol was successful in obtaining genome-wide SNP information for *R. ecuadoriensis*. Sequencing of non-target species was minimal (0.2%) (Supplementary Figure 1). We genotyped six *Rhodnius prolixus* as controls and 80% of reads mapped to the *R. prolixus* reference genome. Only 9.5% of *R. ecuadoriensis* reads mapped to the same reference, a consequence of genomic sequence divergence between *R. ecuadoriensis* and *R. prolixus* ^37^. A stringent genotyping approach confidently identified 2,552 SNP markers across 272 *R. ecuadoriensis* samples from 25 collection sites, which represented closely administrative boundaries of human communities. (Supplementary Table 1). In seven collection sites (Figure 1a; CG, BR, CE, CQ, HY, SJ and GL-seven pairs) triatomines from both domestic and wild ecotopes were collected. Remaining sites only had individuals of one ecotope (domestic or wild).

**Figure 1.**
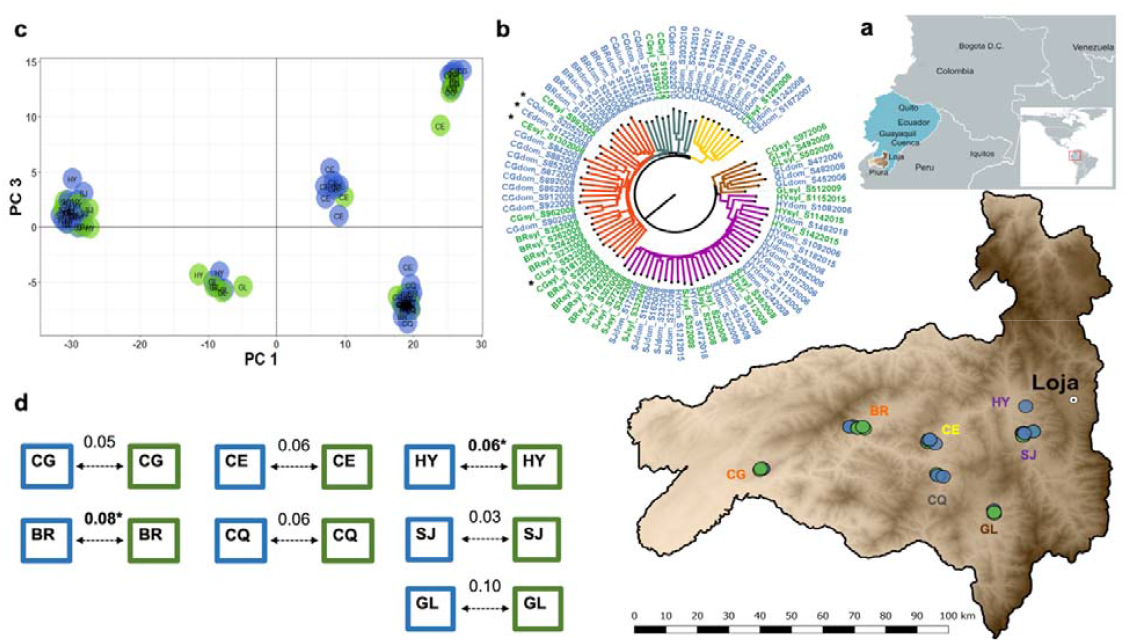
Genomic differentiation of domestic and wild *R. ecuadoriensis*. **a**, geographic distribution of the seven collection sites with both ecotopes over an elevation surface map of Loja. **b**, Neighbor-Joining midpoint phylogenetic tree with phylogenies indicating the Euclidean distance between triatomine samples built from allele counts. Tree branches clades are colour-coded to approximately differentiate collection sites (or clusters of collection sites) with few samples (black asterisks) not conforming to the pattern. **c**, the scatter plot shows five clusters are built with the first and third principal components of the discriminant analysis eigenvalues. **d**, pairwise F_ST_ comparisons between domestic (blue box) and wild (green box) *R. ecuadoriensis* in multiple sites across Loja (**a**). Significant F_ST_ values (arrows) after FDR correction are highlighted in bold and an asterisk. In all panels, samples location (dots) and labels are colour-coded to indicate their domestic (blue) or wild (green) collection ecotope. Collection sites 2-letter ID labels: SJ, San Jacinto; HY, EL Huayco; GL, Galapagos; CQ, Chaquizhca; CE, Coamine; BR, Bramaderos; CG, La Cienega (see Supplementary Table 1 for full collection sites list).

### Reduced *R. ecuadoriensis* population genetic diversity in domestic ecotopes

Multiple genetic diversity estimates among populations from the 25 collection sites in Loja province were calculated (Obsvered (H_O_), and expected (H_E_) heterozygosity, inbreeding coefficient (F_IS_) and Allelic Richness (A_r_); Supplementary Table 2). Sample-size corrected A_r_ values ranged from 1.19 to 1.44 with the lowest values in La Extensa (EX), San Jacinto (SJ), El Huayco (HY) and Santa Rita (RT). In the paired ecotopes within the seven collection sites, A_r_ values were higher for wild than domestic triatomine populations in five out of seven instances, a significant effect observed (p<0.05, rarefaction method^38^).

### Genomic differentiation between domestic and wild ecotopes

To assay populations dynamics between sympatric domestic and wild foci, we focused our individual-based genomic differentiation and pairwise F_ST_ comparisons analyses on the seven collection sites for which samples from both ecotopes were available (Figure 1a). Supporting frequent migration between domestic and wild ecotopes, samples from each ecotope were interleaved at most collection sites in the phylogenetic tree, with collection site geography, not ecotope, impacting the tree topology (Figure 1b). As such, samples collected in Galapagos (GL), Coamine (CE) and Chaquizhca (CQ) formed distinct clusters, and El Huayco (HY) - San Jacinto (SJ) and Bramaderos (BR) - La Cienega (CG) also grouped discretely. Five broadly congruent clusters were defined in a discriminant analysis of principal components (DAPC) (Figure 1c), with geographic collection site rather than their ecotope again structuring observed diversity. F_ST_ indices between paired domestic and wild triatomine samples within each of the seven compared collection sites indicate little differentiation (e.g., F_ST_ ≤ 0.10). Permutation tests indicated that F_ST_ was significant (p < 0.05) at only two sites - Bramaderos and El Huayco (Figure 1d). As expected, hierarchical analysis of molecular variance revealed genetic subdivision was significantly stronger (F_collection sites/total_ = 0.26, p-value < 0.001) among collection sites than among ecotopes within collection sites (F_ecotope/collection site_ = - 0.004, p-value < 0.001) or among collection year within communities (F_collection year/collection site_ = 0.06, p-value < 0.001) (Supplementary Table 4).

### Genetic loci correlated with domestic colonisation

To identify loci associated with domestic colonisation, we combined a Random Forest (RF) classification approach and redundancy analyses (RDA) with outlier scans. We included the seven collection sites with frequent domestic-wild migration and three additional wild-only sites to roughly conform similar number of domestic (n= 56) and wild (n= 52) samples. A total of 347 SNPs provided high ranked classification accuracy (mean > 3) across the three RF iterations (inset in Figure 2a). Backwards purging on this highly discriminatory subset of SNPs detected a set of 43 SNPs that minimised the ‘Out-of-bag’ error rate (OOB-ER) to a minimum of 0.09 and maximised the discriminatory power among domestic and wild samples (Figure 2a). In a parallel RDA model, ecotope (domestic / wild) was a predictor explaining approximately 0.4% of the total variation and the constrained axis built from that variation was significant (p-value < 0.001), and so was the full model as indicated by the Monte Carlo permutation test. The distribution of each SNP loading/contribution to the RDA significant axis showed 109 candidate adaptive loci as SNPs loadings at ±2 SD from the mean of this distribution (permissive threshold; Figure 2b). In a more conservative approach, we also identified seven loci from those 109 under very strong selection as represented by those SNPs loading at the extreme ±3 SD (conservative threshold) away from the mean distribution of the constrained axis (Figure 2b). The arrangement of the individual samples in the ordination space with relation to the RDA axis showed a clear pattern of subdivision comparable to the ecotope in which samples were collected (Figure 2c). The 21 loci/SNPs identified as adaptive loci (dark dots in Figure 2b) by RDA were also detected as highly discriminatory SNPs for domestic and wild ecotopes in the RF analysis. Assuming ‘adaptation with geneflow’ we assessed locus-specific estimates of F_ST_ (Figure 2d), among the 2552 SNPs between domestic and wild ecotopes and identified one SNP (Locus ID 15732 – purple diamond in Figure 2bd) likely to be under local adaptation and/or spatial heterogeneous selection as suggested by OutFlank analysis (Figure 2d left). Moreover, outlier scan with fsthet (Figure 2d right) in the same subset flagged this OutFlank SNP and 73 additional SNPs showing F_ST_ higher that the average neutral loci distribution at a 5% threshold. In summary, 43 SNPs were identified with the highest classification accuracy in RF analysis. 21 of those SNPs showed some signal of adaptation (that is, loaded ± 2 SD away from mean distribution of contrained axis) and 4 were identified showing strong signal of adaptation (that is, loaded ± 3 SD away from mean distribution of contrained axis) in RDA analysis. Three of the SNPs flagged as outliers in fsthet analysis were found also being at high classification accuracy in RF analysis. The SNP (Locus ID 15732) likely to be under strong selection as identified by OutFlank analysis, also had a high classification accuracy in RF and, interestingly, it was also identified within the RDA and fsthet SNPs sets under strong signal of selection.

**Figure 2.**
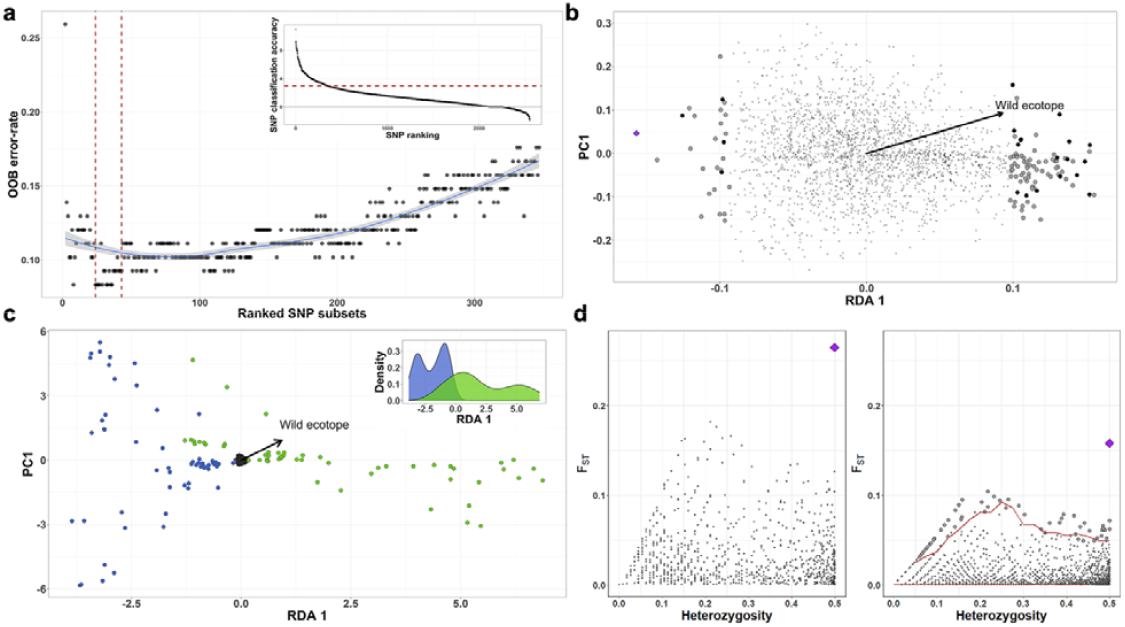
Scanning outlier SNP markers for signatures of local adaptation. **a**, Random Forest backwards purging shows subsets with decreasing number of highly discriminatory SNPs and their resulting OOB-ER. The two vertical red lines indicated the 43 SNPs subset with the lowest OOB-ER and maximum discriminatory power between domestic and wild ecotopes. The inset shows SNPs ranked based on their classification accuracy averaged after 3-independent RF runs. SNPs with classification accuracy above three (red horizontal line) were used for the backwards purging. **b**, In our RDA model, SNPs (dots and diamonds) are arranged as a function of their relationship with the constrained predictor, ecotope (arrow outlines towards a wild ecotope relationship). SNPs closer to the centre (small grey dots) are not showing relation with the predictor. Adaptive loci/SNPs are represented by those large dots/diamond loading at ± 2 SD and ± 3 SD separated from the mean SNPs loading distribution. Black large dots (and purple diamond) represent loci/SNP identified with high classification power in RF analysis. **c**, a biplot of *R. ecuadoriensis* triatomine smaples and SNPs (small black dots in the centre) are arranged in relation to the constrained RDA axis with an arrow indicating those related to the wild ecotope. Dots are colour-coded to show sample ecotope of collection, domestic (blue) or wild (green). Biplot scaling is symmetrical with inset showing the density function for the RDA axis. **d**, Scatter plots show OutFlank (left) and fsthet (right) SNPs F_ST_- heterozygosity relationship. 43 SNPs (large dots) had higher than average F_ST_ distribution of neutral loci in fsthet, whereas only one in OutFlank. Purple diamond indicated the SNP (ID 15732) flagged in all four analyses.

### Mapping outlier loci to the *Rhodnius prolixus* genome

Several SNPs from the different analyses mapped to annotated regions of the *R. prolixus* genome. One SNP identified in the RDA analysis mapped (97.1% identity) in a *R. prolixus* genome region containing the characterised *Krüppel* gap gene (Accession No JN092576.1) involved in embryo development in arthropods^39^. Three SNPs likely to be under balancing selection identified in fsthet analysis mapped (100% identity) to regions in the *R. prolixus* genome containing characterised GE-rich and polylysine protein precursors (mRNA - Accession AY340265.1), and the *Krüppel* and giant gap genes^39,40^ (Accession No HQ853222.1). The former are important proteins within the sialome of blood-sucking bugs^41^ and the latter involved in embryo development^40^. Mapping of the majority of putatively adaptive SNPs, including Locus ID 15732, was not possible in the absence of an available *R. ecuadoriensis* genome.

### Comparison of dispersal rates of *R. ecuadoriensis* between domestic sites with dispersal rates between wild sites

Including all samples (n = 272) and collection sites (n = 25), we tested the strength of genetic isolation-by-distance (IBD) initially among domestic sample collection sites and latterly among wild collection sites (Figure 3). Mantel tests in both domestic (r_m_ = 0.46, p-value < 0.001) and wild (r_m_ = 0.31, p-value = 0.043) ecotopes strongly supported an effect of geographic distance on genetic distance (Figure 3a). Based on a generalised least square model (Supplementary Table 5) with maximum likelihood population effects parametrisation (GLS-MLPE), the effect of geographic distance significantly stronger (0.0018, p-value < 0.001) in wild compared to domestic foci (Figure 3a), suggesting that the rate of vector dispersal occurred at a higher rate between domestic populations than between wild ones.

**Figure 3.**
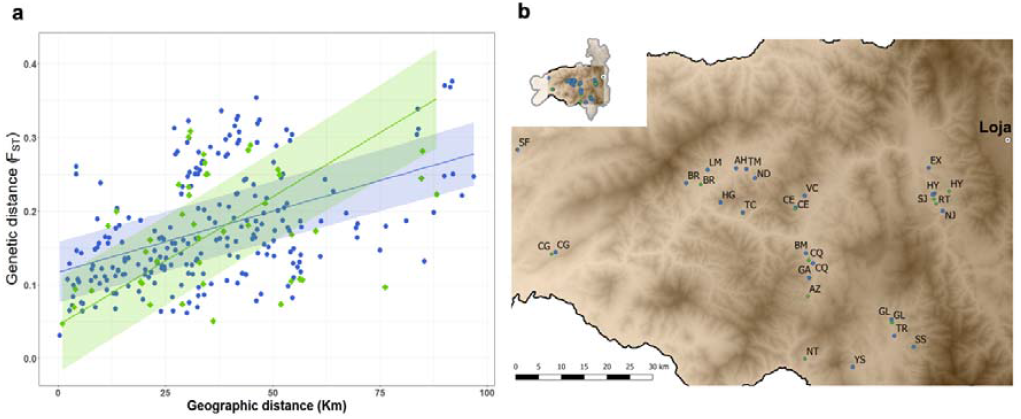
Dispersal rate in *R. ecuadoriensis*. **a**, correlation between pairwise genetic (F_ST_) and geographic distances (data points) with fitted regression lines (95% CI) for domestic (blue dots) and wild (green diamonds) ecotopes. Fitted GLS-MLPE model in eqn 1. **b**, geographic distribution of the 25 collection sites across Loja province used for estimating *R. ecuadoriensis* gene flow with geographic distance. Collection sites 2-letter ID labels: EX, La Extensa; SJ, San Jacinto; HY, EL Huayco; RT, Santa Rita; NJ, Naranjillo; GL, Galapagos; SS, Santa Rosa; TR, Tuburo; YS, Camayos; NT, San Antonio de Taparuca; AZ, Ardanza; GA, Guara; CQ, Chaquizhca; BM, Bella Maria; CE, Coamine; VC, Vega del Carmen; TM, Tamarindo; HG, Higida; ND, Naranjo Dulce; TC, Tacoranga; AH, Ashimingo; LM, Limones; BR, Bramaderos; CG, La Cienega; SF, San Francisco (SF).

### Landscape functional connectivity in *R. ecuadoriensis*

Landscape genomic mixed modellling aims to identify the effect of different combinations of landscape surfaces and their parameters on a given genomic differentiation pattern. To obtain an accurate representation of the genomic differentiation pattern among *R. ecuadoriensis* populations, we chose Hedrick’s G_ST_ pairwise comparisons (Figure 4b) which corrects for sampling limited number of populations^42^. The genomic pattern was consistent regardless of metric used (e.g., Pairwise F_ST_ ^43^ and Meirman’s standardised F_ST_ ^44^) as revealed by strong and significant (r^2^ = 0.99 & 0.92, respectively; p < 0.001) Pearson’s correlations. Pairwise Hedrick’s G_ST_ comparisons showed a strong pattern of population structure across Loja province with presence of both high and low genetic differentiation among collection sites (Figure 4ab). San Francisco (SF) and San Antonio (NT) were two examples of clear, and mutually distinct, outliers in genetic terms. Santa Rita (RT), El Huayco (HY), San Jacinto (SJ) and La Extensa (EX) were genetically and geographically close but highly differentiated form the rest. Overall, clusters of collection sites were evident with some differentiation within and among clusters (Figure 4b).

**Figure 4.**
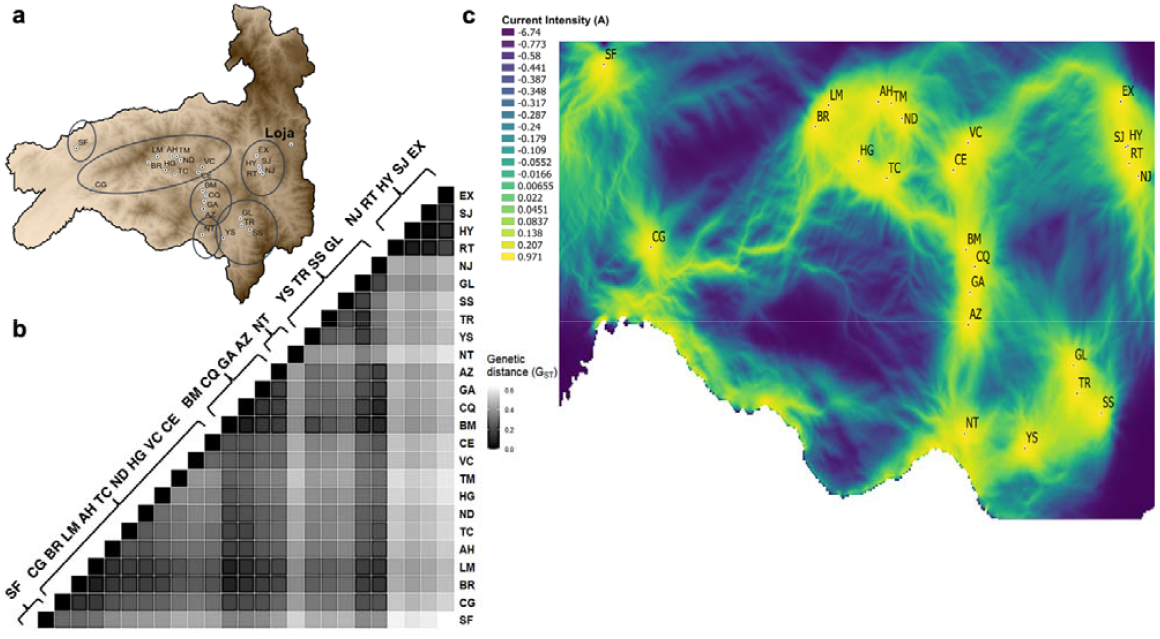
Landscape connectivity of *Rhodnius ecuadoriensis* in Loja province, Ecuador. **a**, Map of the geographic location of collection sites across Loja. **b**, Heatmap shows pairwise genetic distances (G_ST_) with collection sites ID labels on the right. Clusters and highly differentiated collection sites are circled in **a**. Grey scale indicate genetic distance with lighter colours showing higher differentiation. **c**, Electrical current map of Loja built from the optimised elevation surface model showing a gradient of high (yellow/light shade), medium (light greens) and low (blue/dark shade) functional connectivity across Loja. Clusters of highly connected sites are evident but isolated sites are also present across regions on Loja. Connectivity within and among clusters and collection sites is highly influenced by the landscape, specifically elevation surface.

The genomic pattern was iteratively regressed with different combinations of landscape variables and parameters using the ResistanceGA^45^ optimisation framework (see Methods). The optimisation process involves estimating unbiased resistance values for a given combination of surfaces and select the best (true) model representing the genomic pattern. To rule out collinearity between landscape variables, we calculated Spearman’s correlation coefficient, rho, between all pairs of surfaces which resulted in small and/or negative (rho < 0.29) correlations (Supplementary Table 8). Similarly, a scatterplot matrix did not show highly correlated surfaces (Supplementary Figure 11).

Our three ResistanceGA optimisation replicates (see Methods) showed comparable results. In all replicates, the single elevation surface showed the lowest AIC_c_ values and the highest AIC_c_ weight compared to the other single and composite optimised surfaces (Table 1 is a replicate example). Delta AIC_c_ shows the AIC_c_ difference between the elevation surface (best model) and the rest of the (combination of) surfaces. A difference of ∼2.26 units between elevation surface and a distance-only model was evident which suggests elevation surface is a better predictor that geographic distance. Optimisation of the elevation surface parameters confirmed gene flow resistance increases with altitude up to the highest resistance at approximately 2,400 m.a.s.l. (Supplementary Figure 12).

**Table 1.** Model selection results for the generalised mixed-effects models optimised on genetic distance (Hedrick’s G_ST_) for *R. ecuadoriensis*. For each resistance surface model, number of parameters plus the intercept (k), Akaike information criterion (AIC), additional parameters corrected AIC (AIC_c_), marginal (R^2^m) and conditional (R^2^c) R^2^ values of the fitted MLPE model, log-likelihood (LL), delta AIC_c_ and AIC_c_ weight (ω) are provided.

| Resistance surface model | Type | $k$ | AIC | $AIC_c$ | $R^2_m$ | $R^2_c$ | LL | Delta $AIC_c$ | $\omega$ |
| --- | --- | --- | --- | --- | --- | --- | --- | --- | --- |
| Elevation | single | 4 | -751.51 | -749.51 | 0.41 | 0.72 | 379.76 | 0 | 0.76 |
| Distance | single | 2 | -743.79 | -747.25 | 0.43 | 0.74 | 375.90 | 2.26 | 0.24 |
| Roads | single | 6 | -744.91 | -736.25 | 0.42 | 0.75 | 376.46 | 13.26 | 0.0010 |
| Elevation + Roads | composite | 9 | -751.55 | -729.55 | 0.42 | 0.72 | 379.77 | 19.96 | 3.49e-05 |
| Land | Single | 12 | -762.26 | -720.26 | 0.52 | 0.81 | 385.13 | 29.25 | 3.35e-07 |
| Elevation + Land cover | composite | 15 | -763.15 | -687.82 | 0.57 | 0.78 | 385.57 | 61.70 | 3.03e-14 |
| Land cover + Roads | composite | 17 | -762.06 | -648.63 | 0.56 | 0.78 | 385.03 | 100.88 | 9.37e-23 |
| Null model | single | 1 | -561.25 | -565.08 | 0 | 0.42 | 283.63 | 184.43 | 6.75e-41 |
| Elevation + Land cover + Roads | composite | 20 | -762.44 | -520.44 | 0.55 | 0.78 | 385.22 | 229.08 | 1.36e-50 |

To evaluate the roboustness of our optimisation procedure and test the effect of uneven distribution of sample sites, we ran a bootstrap analysis with resampling of the sites at each iteration. Interestingly, the bootstrap analysis revealed that, when resampling 85% of the collection sites, the optimised elevation surface model was ranked the top model in only 43.2% of the bootstrap iterations compared to 46% of the times in which a distance-only model was better (Table 2). The fact that elevation surface was slightly less supported in the bootstrap analysis is likely due to the irregular distribution of sites across the study area and altitudes. Nevertheless, elevation surface remains the strongest predictor of genetic connectivity across the study area (Table 1).

**Table 2.** Summary of bootstrap analysis. For each resistance surface model, number of parameters plus the intercept (*k*), and average (Avg) of the Akaike information criterion (AIC), additional parameters corrected AIC (AIC_c_), AIC_c_ weight (ω), rank, R^2^m, LL, root mean square error (RMSE) and frequency the model was top ranked are provided.

| Resistance surface model | $k$ | Avg AIC | Avg $AIC_c$ | $\omega$ | Avg rank | Avg $R^2_m$ | Avg LL | Avg RMSE | Top model (%) |
| --- | --- | --- | --- | --- | --- | --- | --- | --- | --- |
| Elevation | 4 | -535.59 | -533.09 | 0.40 | 1.62 | 0.41 | 271.79 | 0.055 | 43.2 |
| **Distance** | 2 | -531.44 | -530.78 | 0.60 | 2.33 | 0.42 | 267.72 | 0.055 | 46 |
| **Land cover** | 12 | -526.59 | -487.59 | 4.17e-05 | 3.91 | 0.51 | 275.30 | 0.053 | 10.8 |
| **Roads** | 6 | -523.81 | -517.81 | 0.0008 | 4.18 | 0.41 | 267.91 | 0.055 | 0 |
| **Elevation + Roads** | 9 | -525.76 | -509.40 | 2.80e-06 | 4.40 | 0.41 | 271.88 | 0.055 | 0 |
| **Elevation + Land cover** | 15 | -521.94 | -425.94 | 1.45e-21 | 5.29 | 0.55 | 275.97 | 0.054 | 0 |
| **Land cover + Roads** | 17 | -516.51 | -312.51 | 3.97e-46 | 6.59 | 0.54 | 275.26 | 0.054 | 0 |
| **Elevation + Land cover + Roads** | 20 | -511.14 | 328.86 | 1.11e-185 | 7.68 | 0.54 | 275.57 | 0.054 | 0 |

To assist with the identification of vector management zones for regional health authorities, an electrical current map was built by applying a circuit theory algorithm^46,47^ on the optimised elevation surface model (Figure 4c). Specifically, the algorithm simulates the passing of an electric current across grids (zones) with low/high optimised resistance values. Low resistance grids are highlighted as high current intensity zones (yellow/light zones in Figure 4c) in which high population connectivity, and therefore high degree of gene flow, is predicted. The map showed different gradients of connectivity within and among western, central, eastern and southern Loja province. These included individually isolated populations (e.g. SF & CG), isolated clusters (e.g EX; SJ; HY; RT; NJ); as well as well-connected hubs (e.g., BR-LM, AH-TM-ND, HG-TC and CE-VC).

## Discussion

In this study we make several core observations: *R. ecuadoriensis* do invade houses from wild populations, genomic signatures of *R. ecuadoriensis* domestication can be functionally mapped, and the landscape drivers of vector dispersal can be identified. Consistent with frequent house invasion, high levels of gene flow between multiple domestic and wild *R. ecuadoriensis* populations were detected by hierarchical analysis. Low and largely non-significant pairwise F_ST_ values, as well as interleaved sample clustering based on phylogenetic and discriminant analyses were also consistent with house invasion. Significantly elevated allelic richness in wild sites by comparison to nearby domestic foci clearly confirmed that dispersal occurred most frequently from wild ecotopes into domestic structures. Genome scans across these parallel domestication events revealed strong evidence of ‘adaptation with geneflow’, with key outlier loci associated with colonisation of human-made domestic structures and, presumably, human blood feeding - several of which mapped to the *R. prolixus* genome. A strong signature of isolation-by-distance (IBD) was observable throughout the dataset, an effect less pronounced between domestic sites than between wild foci. Formal landscape genomic analyses revealed elevation surface as the major barrier to genetic connectivity between populations. Landscape genomic analysis enabled a spatial model of vector connectivity to be elaborated, informing ongoing control efforts in the region and providing a model for mapping the dispersal potential of triatomines and other disease vectors.

Vector control is the mainstay of Chagas Disease control^11^. Widespread wild reservoir hosts, as well as a lack of safe treatment options^48,49^ and associated healthcare infrastructure, mean that transmission cannot be blocked by reducing parasite prevalence in human and animal hosts^50^. Our data indicate that elimination of domesticated *R. ecuadoriensis* in Ecuador will be frustrated by repeated re-invasion from the wild environment. Similar risks to effective control are posed by wild *T. infestans* in the southern cone region^12^, *R. prolixus* in Los Llanos of Colombia and Venezuela^13^ and potentially elsewhere in Latin America where competent vectors are present in the wild environment and nearby domestic locales (e.g., *T. sordida, T. maculata, R. pallescens* and others^14,15^).

Understanding evolutionary processes that underpin the colonisation of the domestic environment by arthropod vectors, and their specialisation to feeding on humans, is required to characterize their vectorial capacity. Hybrid ancestry in *Culex pipiens*, for example, is thought to contribute to the biting preference for humans^51^. Human feeding preference can be rapidly genetically selected for in *Anopheles gambiae*^52^. Specialisation of *Aedes aegypti* on humans, and resultant global outbreaks of dengue, yellow fever, and Chikungunya viruses, may be traceable to the emergence of a differential ligand-sensitivity of the odorant receptor *AaegOr4* in East Africa^2^. In triatomines, the nature of genetic adaptions that have enabled the widespread dispersal of successful lineages are far from clear. *T. infestans*, thought to have originated in the Western Andean region of Bolivia, spread rapidly among human dwellings in the Southern Cone region of South America before its near eradication in the 1990s^10^. Cytogenetic analyses suggest this early expansion was accompanied by a substantial reduction in genome size^53^, but the adaptive significance such a change is not clear. The advantage of the *R. ecuadoriensis* system we describe is that it captures multiple parallel adaptive processes and; therefore, can assist in the identification of common evolutionary features associated with colonisation of the domestic environment. Despite limited genomic coverage, and with no *R. ecuadoriensis* reference genome available, we mapped outlier loci to genes in the *R. prolixus* draft genome, and found they are related to salivary enzyme production^41^, as well as embryonic development^39^. Although these genes may have a role in domestic adaptation in triatomines, genome-wide association studies or quantitative trait locus mapping approaches are necessary to fully reveal the genomic architecture of adaptation to the domestic setting. Nevertheless, these findings motivate us to investigate further putative genes involved in local adaptation to the domestic environment such as blood-feeding^54^, sensory cues and host-seeking behaviour^25,55^, as well as human blood detoxification^54,56^. Recent data from our group in Loja province shows that, without doubt, domestic *R. ecuadoriensis* feed extensively on human blood^57^.

Our analyses identified a strong signal of genetic IBD among *R. ecuadoriensis* populations across our study area. Geographic partitioning at this scale is consistent with limited autonomous dispersal capabilities of triatomines which, are, in the main, poor fliers^58^. Wind-blown dispersal observed in smaller vector species is unlikely in triatomines^59^. Passive dispersal of triatomine vectors alongside the movements of their human hosts, which certainly underpins the successful dispersal of other domesticated vector species, is more likely (e.g., *Aedes spp*. ^60,61^). Lower IBD observed among domestic than wild settings may be consistent with passive dispersal alongside humans. We observed a similar phenomenon among parasite isolates from the same region in a previous study^32^. Nonetheless, our formal exploration of the landscape drivers of vector dispersal did not reveal an important effect of roads, and it is not clear to what extent human dispersal of vectors takes place based on our data alone.

According to our landscape genomic analysis, elevation surface is a key predictor of connectivity/discontinuity among *R. ecuadoriensis* populations. Our machine learning (ML) optimisation procedure provides objective parameterisation of altitude resistance values to *R. ecuadoriensis* gene flow^62^. Based on our landscape model predictions we were able to construct a electric current map (Figure 4c) to assist medical entomologists and policy makers in understanding vector dispersal routes. Current vector control strategies in Loja target a single civic administrative unit (neighbourhood or town) for any given insecticidal intervention^28^. Our data and model suggest this approach may be effective for certain communities (e.g., SF, CG, NT and YS, Figure 4). However, for highly connected hubs (e.g. BM, GA, CQ, AZ), successful longer term triatomine control (e.g., insecticide spraying, house improvement, window nets, etc.) will depend on simultaneous intervention in multiple connected communities.

In Ecuador, as with many other endemic regions in Latin America, efforts to control Chagas disease may be complicated in the long term by substantial wild populations of secondary triatomine vectors^16^. As with many other vector borne diseases, there is also a strong case for the use of integrated vector management (IVM) for Chagas disease, where improvements to housing, education, community engagement, in addition to bed net use and insecticide spraying are all likely to be necessary to achieve sustained control^28,63^. Our data clearly indicate that triatomines do invade houses in Loja and low-lying valleys provide routes for vector dispersal between communities and cost-effective IVM must be underpinned by this understanding of vector population structure. Fortunately, genomic and analytical tools can now furnish much of the detail, although better genomic resources for secondary triatomine vector species are required to reveal the process of vector adaptation to the human host. Targeting secondary vector species must now be a priority for health authorities, as these now represent the most pernicious and persistent barrier to controlling residual Chagas disease transmission.

## Methods

### Sample collection and study area

*Rhodnius ecuadoriensis* triatomine bugs (Supplementary Table 1) were derived from a larger collection in the Center for Research on Health in Latin America (CISeAL) of Pontificia Universidad Católica del Ecuador (PUCE). *Rhodnius prolixus* samples (n=6) were provided by the London School of Hygiene and Tropical Medicine and sequenced as an outgroup, as well as to assist with the decontamination of the of 2b-RAD reads and their mapping to functional regions in the draft *R. prolixus* genome^24^. *R. ecuadoriensis* individuals were collected using the one-hour-man method during field surveys across Loja, Ecuador from 2004 to 2018^28^. The triatomines were collected under Ecuadorian collection permits: N° 002– 07 IC-FAU-DNBAPVS/MA; N° 003–2011-IC-FAU-DLP-MA; N° 006-IC-FAU-DLP-MA-2010; N° 010-IC-FAN-DPEO-MAE; N° 011–2015-IC-INF-VS-DPL-MA; MAE-DNB-CM-2015-0030 and internal mobilization guide N° 001-2018-UPN-VS-DPAL-MAE and N° 017-2018-UPN-VS-DPAL-MAE, All these samples were exported to the University of Glasgow by the scientific export authorization N°70-2018-EXP-CM-FAU-DNB/MA.

A widespread spatial sampling (Supplementary Figures 8, 9 and 10) of ecotopes (e.g., domestic and wild), altitudes (up to 1542.9 m.a.s.l.), vegetation types (e.g., tree/bush forest, cropland, etc.) and sites adjacent to different road infrastructure (e.g., highways, tertiary roads, etc.) was carried out in the study area.

### Genomic DNA extraction and sequencing

Genomic DNA (gDNA) was extracted in 88.2% (502/443) of the samples using a SSNT/Salt precipitation method^64^ previously applied in triatomine bugs^65^. For each sample, gDNA concentration was > 25 ng/uL and 288.4 ng/UL (sd. ± 241.8) on average with purity ratios (260/280 and 260/230) of 1.87 (sd. ± 0.10) and 2.30 (sd. ± 0.97), respectively. gDNA was digested with the CspCI Type IIB restriction enzyme (IIB-REase - New England BioLabs, Inc.) which has shown to yield a high marker density in triatomine^65^. DNA fragments (36bp) were ligated to Illumina single-end adaptors and a specific barcode added during PCR amplification to construct 382 150bp 2bRAD libraries^66^. Libraries were homogenised to an approximate similar concentration, purified with magnetic beads^67^ and pooled in two separate batches (n = 191). Each batch was sequenced separately on 1-flowcell (2 lanes) HiSeq 2500 (Illumina) Rapid Mode platform with a single-end (1×50 bp) setup using v2 SBS chemistry at the Science for Life Laboratory (SciLifeLab, Stockholm, Sweden), which also implemented the reads demultiplexing and their in-house quality-filtering.

### Bioinformatics of 2b-RAD sequenced data

#### Data cleaning and decontamination

Demultiplexed raw data quality scores were verified in FastQC software v0.11.9 (http://www.bioinformatics.babraham.ac.uk/projects/fastqc/). 2.3% (16/689) Million reads (Mreads) were removed due to incomplete CspCI restriction site (36 bp) and having across read quality score below 30^68^. The 624.7 high quality Mreads with integrate restriction site had their Illumina adaptors and barcodes trimmed, and reads were forwarded (5’-3’) using custom scripts. To exclude non-target sequences (Supplementary Methods 1.1), 1.2 Mreads (0.2%) reads mapping to bacteria, virus, archaeal, *Trypanosoma cruzi*^69^ and *homo sapiens* (Genome Reference Consortium human build 38) genomes were removed using DeconSeq standalone v4.3^70^ with an alignment identity threshold of 85% and Kraken^71^ taxonomic classifier (Supplementary Figure 1). After decontamination, each sample yield on average 1.6 Million reads (interquartile range = 1.9 Mreads).

#### Optimisation and genotyping

As advised in refs.^72,73^, we optimised (Supplementary Methods 1.1) STACKS v2.55^74^ DENOVO_MAP.PL programme by varying at a time one of the main controlling parameters (-m, -M and -n; Supplementary Table 2) on each run while keeping the rest of the parameters at the setting used in early experiments (e.g., -m 5, -M 2, -n 1, -N 4, -alpha 0.01, -bound_low 0, -bound_high 0.01, -r 0.8, -min_maf 0.01^65^). The parameter combination yielding the highest number of SNPs with the least missing data and genotyping error rate was chosen to be the optimal set. Genotypes below a quality score of 30, and samples with above 50% missing genotypes across sites and among loci were removed from downstream analysis using the VCFtools software suite v0.1.5 5^75^. The remaining missing genotypes (< 0.5%) were imputed using the k-nearest neighbour genotype imputation (LDkNNi) method^76^ implemented in the TASSEL software v5^77^.

### Genomic differentiation between domestic and wild ecotopes

#### Genetic diversity and linkage disequilibrium

Genetic diversity measures (e.g., observed (H_O_) and expected heterozygosity (H_E_), inbreeding coefficient (F_IS_) and percentage of loci in Hardy-Weinberg equilibrium (% HWE)) were calculated for each collection site, and ecotopes (domestic and wild) within collection sites, in the HIERFSTAT^78^ and pegas^79^ packages in R^80^. Sample-size corrected Allelic richness (Ar) was calculated using the rarefaction method^38^ implemented in the PopGenReport^81^ R package. To evaluate the percentage of SNP markers in linkage disequilibrium (LD), correlation coefficient (r^2^) estimates were calculated between markers pairs using using the GUS-LD R package^82^ which revealed a very low percentage (< 0.20%). To observe whether genetic diversity difference between ecotope pairs was significant, a permutation-based (10,000 permutations) two sample t-test was performed on each pair diversity values using the RVAideMemoire R package (https://www.rdocumentation.org/packages/RVAideMemoire).

#### Individual-based genomic differentiation

Genomic differentiation among *R. ecuadoriensis* domestic and wild samples within a subset of seven collection sites was visualised in a neighbour-joining midpoint tree^83^ (Figure 1b) built from Euclidean genetic distances of allele frequencies with the ape^84^ R package. Tree components were edited in FigTree software v1.4.3 (http://tree.bio.ed.ac.uk/software/figtree/) to better illustrate domestic and wild samples and their overall clustering pattern. To explore samples genomic differentiation further, a DAPC^85^ was performed in the same seven collection sites with the adegenet^86^ R package (Figure 1c). The most likely *a priori* number of clusters was chosen based on the lowest Bayesian information criterion (BIC). In the DAPC, all principal components (PCs) and the eigenvectors of the first three DA discriminant functions were kept for visualizing the samples individual coordinates of different PCs linear combinations (Supplementary Figure 5).

#### Pairwise F_ST_ comparisons

To support previous hierarchical analyses, pairwise F_ST_ comparisons^43^ were performed between *R. ecuadoriensis* from domestic and wild ecotopes within the seven collection sites (Figure 3b). In this study, F_ST_ was exploited as a measure of genomic connectivity (flow) between ecotopes within given collection sites. Specifically, Nei’s F_ST_^87^ pairwise comparisons were computed in adegenet R package and tested at 5% significance via 999 permutations of individuals selected randomly within and between groups. P-values were corrected for multiple comparisons using the false discovery rate (FDR) method^88^ in the function p.adjust of the stats R package^80^.

#### Hierarchical F-statistics

*R. ecuadoriensis* molecular variation was explored at a four-level (e.g., among collection sites, among ecotopes (domestic or wild) within collection sites, among collection year within collection sites and among individuals within populations) hierarchy of population structure. For each hierarchy, a F-statistic (with 95% C.I.) was calculated, and its significance tested via 999 randomised permutations with the HIERFSTAT R package. For comparison and given not all sites had both ecotopes, two hierarchical analysis were performed, one with the total collection sites (n = 25) and the other with a subset of collection sites (n = 7) with samples collected in both ecotopes (Supplementary Table 4).

### Domestic-wild SNP association analyses

As a response of *R. ecuadoriensis* ecotopes fluxes in multiple collection sites across Loja, we screened for SNP RADseq markers under a strong signal of selection (outlier loci). The power for detecting outlier loci of four different approaches, Random Forest (RF) machine learning (ML) classification algorithm (implemented in refs.^89–91^), redundancy analysis (RDA) constraint ordination^92^, and OutFlank^93^ and fsthet^94^ F_ST_-outlier methods, was evaluated using a roughly similar number of domestic (n= 56) and wild (n= 52) *R. ecuadoriensis* across Loja province sharing a total of 2552 SNPs.

#### Random Forest

The RF algorithm^95^ implemented in the randomForest^96^ R package was used to build a series of recursive decision trees, or forest, to classify domestic and wild *R. ecuadoriensis* based on their shared SNPs (predictors) covarying to a specific ecotope (response variable) (Supplementary Figure 6). Within each RF run, decision trees were trained by random subsampling with replacement 66.6% of triatomine samples (training dataset), for which aleatory selected SNPs were top-ranked classifiers when minimizing the most within-ecotope variation (that is, partitioning triatomine by ecotope). Trained trees predictive power was tested with the remaining 33.3% triatomine samples (‘Out-of-bag’ test dataset) in which ecotope misclassification of samples estimated an OOB-ER for that RF run; SNPs importance classification accuracy was averaged among the total number of trees created in a given RF. Three independent (spatial structure-corrected) RFs with 100,000 trees were run and their convergence on SNPs importance classification accuracy was evaluated by Pearson’s correlation test. Top-ranked SNPs (Figure 2a inset) among the three RFs (that is, importance classification accuracy above 3) were chosen for backwards purging, as implemented in refs.^90,97^. Backwards purging (Figure 2a) iteratively runs RFs starting with the full top-ranked SNPs and discarding the least important ones before the next iteration until only two were left. The subset with the lowest OOB-ER contained SNPs outlying strongly for the ecotope response.

#### Redundancy analysis

Outlier loci likely under selection were also identified using RDA multivariate constrained ordination^98^ implemented in the vegan^99,100^ R package. First, a matrix fitted values (Supplementary Figure 7a) were obtained using multivariate linear regression between a matrix of genotypes (response) and ecotopes (explanatory) with an additional term controlling for spatial structure (based on the three first axes of an individual principal coordinates of each sample). Then, principal component analysis (PCA) on the fitted values matrix resulted in a constrained axis composed from the variation explained, ‘redundancy’, by our explanatory variable (Supplementary Figure 7b). Overall RDA model and variation explained by the constrained RDA axis were tested for significance via 999 permutations designed for constrained correspondence analysis. Additionally, SNPs coordinates were scaled and plotted in the ordination space to see their relationship with the constraint axis, ecotope (Supplementary Figure 7c and Figure 2b). SNPs z-transformed loadings (Supplementary Figure 7d) separated by ±2 and ±3 standard deviations (permissible and conservative thresholds, respectively) from the mean distribution of the total SNPs loadings in our RDA axis were considered under selection (Figure 2b) (for further details on RDA see refs.^92,101,102^).

#### F_ST_-Heterozygosity outlier method

The F_ST_-Heterozygosity outlier method aims to identify loci with strong allele differences among ecotopes. First, ecotope differentiation for each locus is calculated using Wright’s F_ST_ without sample correction. The distribution of these values is expected to have a chi-squared shape. The main goal is inferring a null F_ST_ distribution from neutral loci not strongly affected by diversifying selection^93^. Therefore, a best-fit to the chi-squared F_ST_ distribution was achieved by trimming the lowest and highest F_ST_ values (loci in the tails of the distribution are likely to be under effective diversifying selection) and considering only the values in the centre (neutral loci and loci experiencing spatial uniform balancing selection). Loci with unusual F_ST_ values relative to this fitted distribution can be thought of experiencing additional diversifying selection^93,94^. We used two R packages to accomplish this analysis, OutFlank^93^ and fsthet^94^, and compared the results (Figure 2d). The difference between the packages is that fsthet uses smoothed quantiles of the empirical F_ST_-Heterozygosity distribution to identify outlier loci and does not assume a particular distribution or model of evolution as compared to OutFlank. We set OutFlank function with proportion of lower and upper loci trimmed to 0.06 and 0.35, respectively, and the rest of the values to default.

#### Mapping SNP outlier loci

In order to identify genes that may be responsible for local adaptation in the Chagas disease vector, *R. ecuadoriensis*, to the domestic environment we mapped the SNPs found in the association analyses to the *R. prolixus* annotated genome^24^. We used the BWA alignment tool implemented in DeconSeq software v0.4.3^70^ to map SNPs sequences (38 bp) at a minimum alignment threshold of 85. The sequences of the regions (60-300kb) in which our SNPs aligned were BLAST searched and compared to the *R. prolixus* genome.

#### Estimating gene flow with distance

Matrices of genetic (F_ST_^87^) and geographic (Km) distances between the 25 collection sites, and between domestic and wild collection sites separately, were obtained with the adegenet and raster^103^ R packages, respectively. Mantel tests^104^ were performed on those matrices using the ecodist^105^ R packages. Genetic and geographic correlation between domestic and wild ecotopes was also viewed separately by fitting a generalised least square (GLS) model with a maximum likelihood population effects correction (MLPE)^106^ implemented in the corMLPE (https://github.com/nspope/corMLPE/) R package and assuming a linear relationship

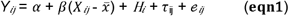

between two distance matrices based on genetic and geographic distance measures, Y and X, respectively. Centring the *X*_*ij*_ in about its mean, 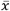, removes the correlation between the estimates of *α* and *β*^106^. H, determines the ecotope and the τ _ij_ term adds the MLPE random effect correlation structure.

### Estimating gene flow with resistance

#### Genetic distances

Given genomic differentiation between domestic and wild ecotopes was low, we combined all samples within a collection site and used collection site as the unit in our landscape genetic analysis. Collection site units are logistically and budgetary important when carrying out triatomine surveys and insecticide spraying. Using a landscape genomics mixed modelling framework (Supplementary Figure 2), we aimed to disentangle the effects of landscape heterogeneity on *R. ecuadoriensis* population structure and gene flow. A Hedrick’s G_ST_ ^42^, which corrects for sampling limited populations^107^, distance matrix among the 25 collection sites was obtained in the GenoDive v3.04^108^ software. In addition, we ran a Pearson’s correlation test between the Hedrick’s G_ST_ matrix, and Meriman’s standardised F_ST_^44^ and F_ST_^43^ matrices, calculated in the same software, to evaluate the consistency of genomic differentiation pattern among collection sites with different genetic distance measures.

#### GIS data collection and preparation

Three landscape variables (elevation, land cover and road network - hereafter, surfaces) were hypothesized to influence *R. ecuadoriensis* dispersal and gene flow (Supplementary Figures 8, 9 and 10). For the continuous surfaces (elevation surface), only monomolecular transformations (e.g., Supplementary Figure 12ab) with any possible shape and maximum parameters were explored to assume a linear relationship in which gene flow decreases as altitude increases as hypothesised in other triatomine species^109^. Our categorical surfaces, land cover and road network, were reclassified as follows. Highly fragmented land cover categories (e.g., cultivated and managed areas) produced the least resistance to gene flow, whereas regular flooded areas and water bodies were barriers. Habitat fragmentation and human agricultural activities has been shown to affect triatomine populations dynamics^110^. Human-mediated passive triatomine dispersal has been suggested elsewhere^11^, and therefore, we assumed roads would connect humans populations, and likely triatomines by passive carriage. High transitable roads (e.g., highways and tertiary roads) had the least resistant values, whereas absence of roads was a strong barrier (see Supplementary Table 9 & Supplementary Table 10). Original GIS surfaces were obtained from multiple sources (Supplementary Table 6) and transformed to have the same format (raster), resolution (250 m^2^ grid), extent (∼ 97 Km^2^) and coordinate reference system (Universal Transverse Mercator (UTM)). Spearman’s rank correlation coefficient (rho) tests were run (Supplementary Table 8) and plotted (Supplementary Figure 11) on each pair of surfaces to ensure variables were uncorrelated (rho < 0.29 based on Cohen^111^). All three surfaces original values were transformed to the same scale (i.e., a minimum value of 1 and a maximum of 100) to meet our initial hypothesis.

#### ResistanceGA principle

The genetic algorithm^112^ implemented in the R package, ResistanceGA^45^, was used for multiple and sinlge-surface optimization of resistance values to gene flow in the above surfaces (Supplementary Figure 3). Briefly, ResistanceGA method is a powerful, flexible, stochastic and assumption-free framework based on an evolutionary ML process that finds unbiased optimal resistance parameters (resistance weights values for a given surface) in space that best fits the genomic structure pattern^113^. The method works by correlating genomic (response) and effective (predictor) distances (derived from a random-walk commute time algorithm^114^ – Supplementary Figure 4b) matrices through a maximum likelihood population effects^106^ model and, on each iteration, evaluates the best resistance parameters based on a ML objective function, log-likelihood in our case. Simulating the process of evolution on each iteration, the best model and parameters are selected and pass over the next generation with some random change on parameter values to explore the parameter space widely.

#### Multiple surface optimization

We performed three replicate runs to optimise all possible combinations of our surfaces (hereafter, composite surfaces), including surfaces individually (hereafter, single surfaces) to generate models with optimised resistance values. The major GA algorithm options were set to default, except for the ‘pop.mult’ which was set to 20 to increase the number of parameters to evaluate on each surface every iteration. All optimisation processes were run in parallel with 10-20 cores in a Debian cluster (http://userweb.eng.gla.ac.uk/umer.ijaz/#orion) at the University of Glasgow. Running times varied from days to weeks depending on surface size and number combined at a time.

#### Model selection

Composite and single surface models, including an intercept-only (null model) and a geographic distance (resistance grid cells are set to 1 to model isolation-by-distance) model were evaluated (Table 1) and the best model was selected based on the lowest AIC_c_, AIC_c_ weight and Delta AIC_c_. To confirm the robustness of the optimization surfaces and controlling for potential bias due to uneven distribution of sample locations in the landscape, we carried out bootstrap resampling (10,000 iterations) in 85% of our sample locations and then fit the subset to each of the effective distance matrices from the optimised surfaces. After the bootstrapping analysis, the average AIC_c_ among all iterations and the percentage a model was top over all iterations was used as a criterion to rank the best model (Table 2).

#### Landscape connectivity model

We used the best optimised single (elevation surface) resistance surface models to estimate landscape connectivity through a circuit theory algorithm^46,47^ (Supplementary Figure 4) implemented in the software CIRCUITSCAPE v5^115^. Here, our resistance surfaces were converted into electric networks (Supplementary Figure 4c) in which each grid cell represented a node connected to their neighbours by resistors of different weight. Resistor weights were calculated from the average resistance values (i.e., optimised resistance values) of the two grid cells being connected. The algorithm applies a simulated electric current between all pairs of focal nodes (collection sites) in the network to estimate effective distances between them. A current density map (Figure 4c) was obtained from those resistance distance estimations representing a random walk probability of movement through our study area.

## Data availability

Raw sequenced data will be uploaded to the Sequence Read Archive (SRA) repository on publication.

## Meterials & Correspondance

Correspondance to Martin Llewellyn and Luis Enrique Hernandez Castro

## Code availability

Code for population, assosiation and landscape genomics analyses will be available via Github repository (github.com/lehernandezc/recuadoriensis) on publication.

## Acknowledgements

We thank P. Johnson for advice in statistical analyses, W. Peterman for helpful advice on ResistanceGA analysis, the entomological team at CISeAL for sample collection and M. Babbucci for proving custom scripts for 2b-RAD raw data cleaning. This work was possible thanks to the Mexican Council of Science and Technology doctorate scholarship (CVU Number 613766) awarded to L.E.H.C., the National Institutes of Health (NIH) grant number R15 AI105749-01A1 allocated to MJG who is PI, as well as RCUK Engagment Network (EP/T003782/1) which supported co-author interactions. Funding was also received from Pontifical Catholic University of Ecuador to MJG (grant # C13025, E13027, E13037, H13174, I13048). ELL was supported by the National Institute of General Medical Sciences of the NIH, United States (Award Numbers P20GM130418). The funders had no role in study design, data collection and analysis, decision to publish, or preparation of the manuscript.

## Author contributions

L.E.H.C., M.S.L., and M.J.G. designed the study. L.E.H.C. and M.S.L. wrote the manuscript with contributions from M.J.G., J.A.C., and E.L.L. A.G.V., A.J., B.C., C.C.D., S.O.M., C.A.Y., A.B., B.A., L.M., E.L.L. revised and edited the manuscript. A.G.V., S.O.M., C.A.Y., and J.A.C. collected and provided triatomine samples from Loja Ecuador. L.E.H.C performed DNA extraction and 2b-RAD library preparation. B.A. performed NGS Illumina HiSeq. L.E.H.C analysed the data with contributions from A.J. in the association analysis, B.C. in data decontamination/bioinformatics, C.C.D., and E.L.L in the landscape genomic analyses.

## Supplementary information

Supplementary Table 1 (excel file)

Supplementary Methods (docx file)

## Figures 1 - 4

Figures in the order of appearance in main text.

