## Supplementary Methods for "The genomic basis of domestic colonisation and dispersal in Chagas disease vectors"

**Supplementary Information**

### 1 **Supplementary Methods**

#### 1.1 **Bioinformatics of 2b-RAD sequenced data.**

**Data decontamination.** Our decontamination pipeline (Supplementary Figure 1) started by mapping *R. ecuadoriensis* reads to the *R. prolixus* annotated genome^1^ using the Burrows- Wheeler algorithm^2^ (BWA) implemented in DeconSeq standalone v4.3^3^. We parametrised the DeconSeq programme by running it on a subset of *R. prolixus* high-quality 2b-RAD reads against the *R. prolixus* reference genome and varying the alignment identity threshold (-i) at 75, 85 and 95. After this trial, the DeconSeq run on the *R. ecuadoriensis* reads was set to 85. *R. ecuadoriensis* reads that mapped to the *R. prolixus* reference genome were kept and unmapped reads were further decontaminated. In the next step, *R. ecuadoriensis* reads that did not map to the *R. prolixus* genome were classified based on a bacterial, archaeal and viral genome database in the Kraken^4^ programme using its taxonomy classification algorithm. *R. ecuadoriensis* reads classified as bacterial, archaeal and viral genomes by Kraken were discarded from the pipeline and those unclassified continued into the next decontamination step. Subsequent decontamination steps involved running again the DeconSeq programme on the unclassified *R. ecuadoriensis* reads, first, against the *T. cruzi* I Sylvio X10/1 genome^5^ obtained from TriTrypDB.org genome database^6^ and, then, against the human reference genome build 38 obtained from the National Centre for Biotechnology Information (NCBI) FTP server (ftp://ftp.ncbi.nih.gov/genomes/H_sapiens/Assembled_chromosomes/seq/).	*R. ecuadoriensis* reads that mapped to the *T. cruzi* and Human genomes were discarded from the pipeline, whereas the rest that did not map were kept. Finally, those *R. ecuadoriensis* reads that did not map to any organisms above were merged with the reads that mapped to the *R. prolixus* genome at the beginning of the pipeline. At the end of the decontamination process, samples below a threshold of 100,000 decontaminated reads (hereafter called clean reads) were not included in the genotyping pipeline.

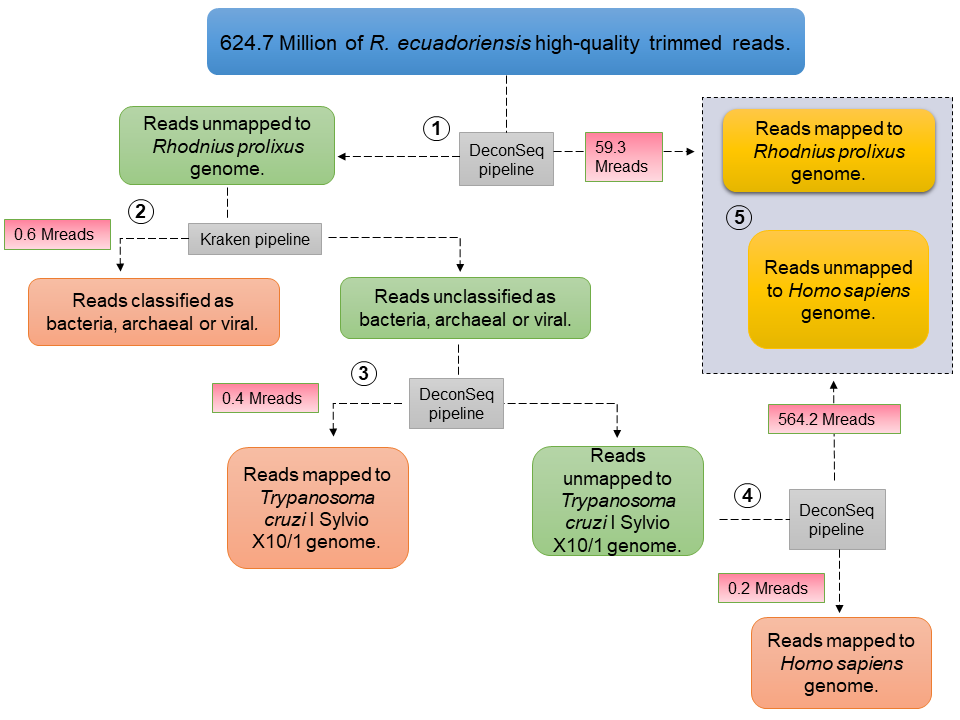

**Supplementary Figure 1 *R. ecuadoriensis* high-quality trimmed reads decontamination pipeline.** Diagram shows the steps carried out to remove contaminant sequences from the *R. ecuadoriensis* high-quality trimmed reads. 1. *R. ecuadoriensis* reads were mapped against *R. prolixus* genome using DeconSeq programme^3^ in which mapped reads (yellow box) were kept and unmapped reads (green box) were further decontaminated. 2. Unmapped reads (green box) were classified based on a bacterium, archaeal and viral database of the Kraken programme (Wood 2014) and the resulting classified reads (red box) were discarded whereas unclassified reads (green box) were passed onto next steps. The “free from bacteria, archaeal and viral” reads (green box) were mapped against *T. cruzi* (3) and human genomes (4) using the DeconSeq programme. In both cases, mapped reads (Red boxes) were discarded from the pipeline and the reads that did not map to neither of genomes (yellow box) were merged (5) to the *R. ecuadoriensis* reads mapped to the *R. prolixus* genome at the beginning of the pipeline (dashed purple box).

**Genotyping optimisation.** The pipeline optimisation consisted in testing eight different combinations (Supplementary Table 2) of the STACKS^7^ DENOVO_MAP.PL programme main control parameters (-m, -M, -n and -N) and selecting the combination resulting in the highest number of polymorphic loci but maintaining a low percentage of missing data and error rate^8,9^. To increase computational speed, we limited genotyping to a representative sample (n = 81) of the complete dataset consisting of 75 *R. ecuadoriensis* samples from six communities (SF, CG, BR, HY, GL, CQ) widely distributed in Loja and one from Manabí, and six *R. prolixus* samples. We used the STACKS parameter combination in our previous *R. ecuadoriensis* genotyping study^10^ as a reference for the first run (Table 3 setting ID m5). In subsequently runs (Table 3 setting ID m4, m3, m2, M3, n0, n2 and n3), we varied one of the main control parameters at a time and kept the reference combination values for the rest of the parameters. Three values of parameter -m (4,3 and 2), one of parameter -M (3) and three of parameter - n (0, 2 and 3) were evaluated. As a reminder, parameters -m and -M are likely to have the most impact on the number of loci yielded. Setting -m too high will drop out true alleles of the pipeline, whereas low values will tend to increase potential sequencing error. Setting -M too low will increase the probability of missing SNPs, whereas low values will allow enough mismatches to build nonsensical biologically incorrect alleles. After -m and M have perfectively assembled loci, then -N recovers a set of “secondary” putative loci that did not achieved enough depth with -m, these set of secondary alleles aid the SNP calling model in detecting polymorphisms as they increase read depth. Finally, -n will look into the population catalog, consensus of all discovered loci, and it will try to merge loci across samples. Setting -n too will allow representation of independent loci in the catalog that in reality are the same locus^7^.

**Supplementary Table 2 Description of the STACKS main control parameters and combinations tested for de novo assembly optimisation.** -m is the minimum number of raw reads required to form a stack (a putative allele) which is comparable to the minimum depth of coverage. - M is Number of mismatches allowed between stacks (putative alleles) to merge them into a putative locus which is comparable to the number of nucleotide mismatches allowed. -n is the number of mismatches allowed between stacks (putative loci) during construction of the catalog that contains all loci and alleles of the population. -N is the number of mismatches allowed to align secondary reads (reads that did not form stacks) to assemble putative loci to increase locus depth. -alpha is the significance level to call a heterozygote or homozygote. -bound_low and -bound_high set the bounded SNP calling model for identifying a SNP and estimating the error rate at that SNP. -r is the percentage of individuals that must possess a particular locus for it to be included in calculation of population-level statistics. -min_maf specify the minor allele frequency for a particular locus, alleles occurring below this frequency are discarded. Further details in refs.^7–10^.

| ***Setting ID*** | ***-m*** | ***-M*** | ***-n*** | ***-N*** | ***-alpha*** | ***-bound_low*** | ***-bound_high*** | ***-r*** | ***-min_maf*** |
| --- | --- | --- | --- | --- | --- | --- | --- | --- | --- |
| *m5** | **5** | **2** | **1** | **4** | **0.01** | **0** | **0.05** | **0.8** | **0.01** |
| *m4* | **4** | 2 | 1 | 4 | 0.01 | 0 | 0.05 | 0.8 | 0.01 |
| *m3* | **3** | 2 | 1 | 4 | 0.01 | 0 | 0.05 | 0.8 | 0.01 |
| *m2* | **2** | 2 | 1 | 4 | 0.01 | 0 | 0.05 | 0.8 | 0.01 |
| *M3* | 5 | **3** | 1 | 5 | 0.01 | 0 | 0.05 | 0.8 | 0.01 |
| *n0* | 5 | 2 | **0** | 4 | 0.01 | 0 | 0.05 | 0.8 | 0.01 |
| *n2* | 5 | 2 | **2** | 4 | 0.01 | 0 | 0.05 | 0.8 | 0.01 |
| *n3* | 5 | 2 | **3** | 4 | 0.01 | 0 | 0.05 | 0.8 | 0.01 |
| * STACKS parameter combination used in ^10^ | | | | | | | | | |

#### 1.2 **Estimating gene flow with resistance.**

**Landscape genetics framework on arthropod vectors.** Landscape genetics^11^ can complement our understanding of arthropod vectors dispersal dynamics by estimating functional connectivity^12^, the level at which the landscape facilitates or impedes their movement from, and to, different habitat patches^13^. This approach involves several stages starting with defining a clear research question which is typically related to investigating landscape arrangement effects on gene flow and/or local adaptation^14^ (Supplementary Figure 2a). At this stage, the extent and resolution of an a priori landscape model is hypothesised based on empirical knowledge of factors (e.g. relief, temperature, land cover etc) likely affecting gene flow of the species of interest. Alternatively, simulation studies could help with specifying a landscape model and supporting spatial and genetic sampling design^15^. Then, pairwise population genetic distance (GD) information (Supplementary Figure 2bc) is obtained by sampling and genotyping individuals (or populations) across the defined landscape model at different gradients and time periods: both spatial and temporal scales are important considerations in a landscape genetic study^14,16^. Subsequently, the hypothetical landscape model is parametrised (Supplementary Figure 2d) in order to obtain a resistance surface (spatial representation of a species movement constraints at each grid cell on a digital layer) from which pairwise population effective distances (ED) estimates of landscape connectivity (comparable to pairwise genetic data) are calculated using least-cost path, commute distance or circuit theory methods^17^. Finally, pairwise population genetic and landscape connectivity distances among points are correlated (Figure 2e) by computing available statistical approaches^18^ such as mantel tests, regression, Bayesian inference, ordination techniques, and more recently, mixed effect models with maximum likelihood population effects parametrisation^19^.

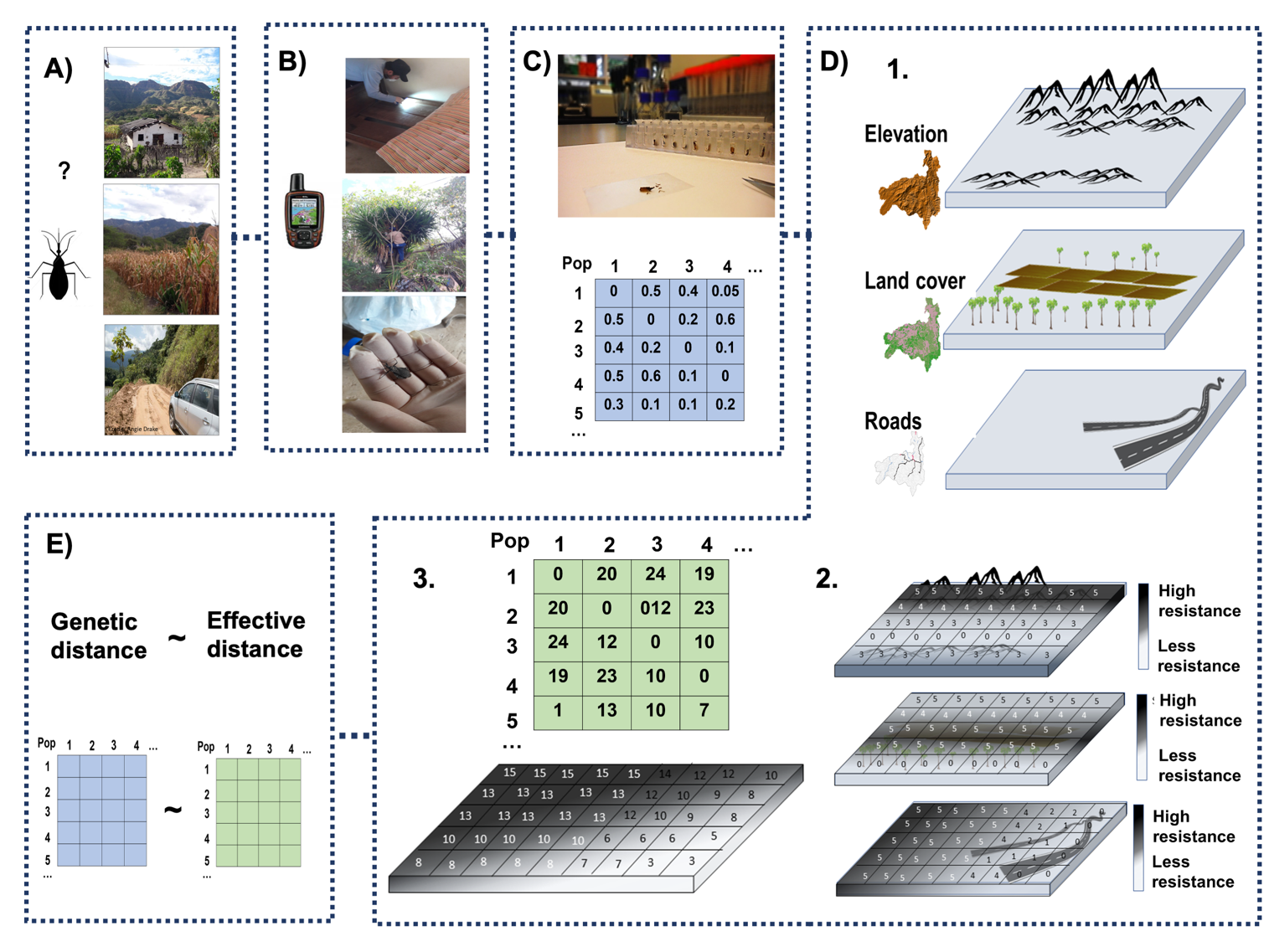

**Supplementary Figure 2 Step-by-step of the landscape genetics framework used to study our Chagas disease arthropod vector, *Rhodnius ecuadoriensis*.** **a**, First, a research question is defined based on whether gene flow or adaptation process are to be investigated and sampling design is stablished. **b**, In the field, triatomine are collected in different ecotopes in the spatial and temporal gradients defined in A). Different variables are recorded at this stage such as altitude and geographic coordinates. **c**, In the laboratory, triatomine next generation sequencing (NGS) libraries are prepared and sequenced in in high-throughput platforms. NGS data is processed and genotype information of each sample is used to obtain a matrix of pairwise populations (Pop) genetic distances. **d**, A hypothetical landscape model (1) is parametrised into a resistance surface (2) which is spatial representation of a species movement constraints at each grid cell on a digital layer. From this resistance surface, a matrix of pairwise population (Pop) effective distances is calculated (3). **e**, Finally, statistical methods are used to correlate matrices of pairwise population genetic (GD) and effective (ED) distances to investigate whether isolation-by-resistance is a fitted model of the genetic differentiation of triatomine populations.

**ResistanceGA optimisation process.** The ResistanceGA algorithm (Supplementary Figure 3) considers an initial population made of each raster surface as individuals experiencing evolution over generations. Raster surfaces resistance weights are the parameters to be optimised (Supplementary Figure 3a). Fitness of a set of parameters is evaluated through an objective function which trains the genetic algorithm (Supplementary Figure 3b). The training is made every time pairwise genetic (response) and effective (predictor) distances are regressed using a linear mixed-effects models with maximum likelihood population effect (MLPE) parametrization. Effective distances are calculated from the individual parameters being evaluated at each iteration (see below details on effective distances calculation). In our case, the machine learning objective function was based on the MLPE model which used the log-likelihood to quantify model performance. In machine learning, the objective function provides a way to assess parameters fitness, the most feasible parameters will have a better chance to be the “parents” of the next generation, but mutation will control stochasticity in the process^20,21^. MLPE regression overcomes the non-independence problem attached to pairwise data^22^ and has been shown to be the best model selection method in landscape genetic studies (see refs.^23,24^). Different combinations of parameters (also seen as “genotypes” within an evolutionary context) in each individual are evaluated iteratively every generation and the fittest individuals (selection) and their parameters are passed over (crossover) into the next generation with top parameters slightly changing at random (mutation). The whole optimisation is repeated until raster surfaces and parameters (model fit) cannot be improved for several generations (convergence check - Supplementary Figure 3c).

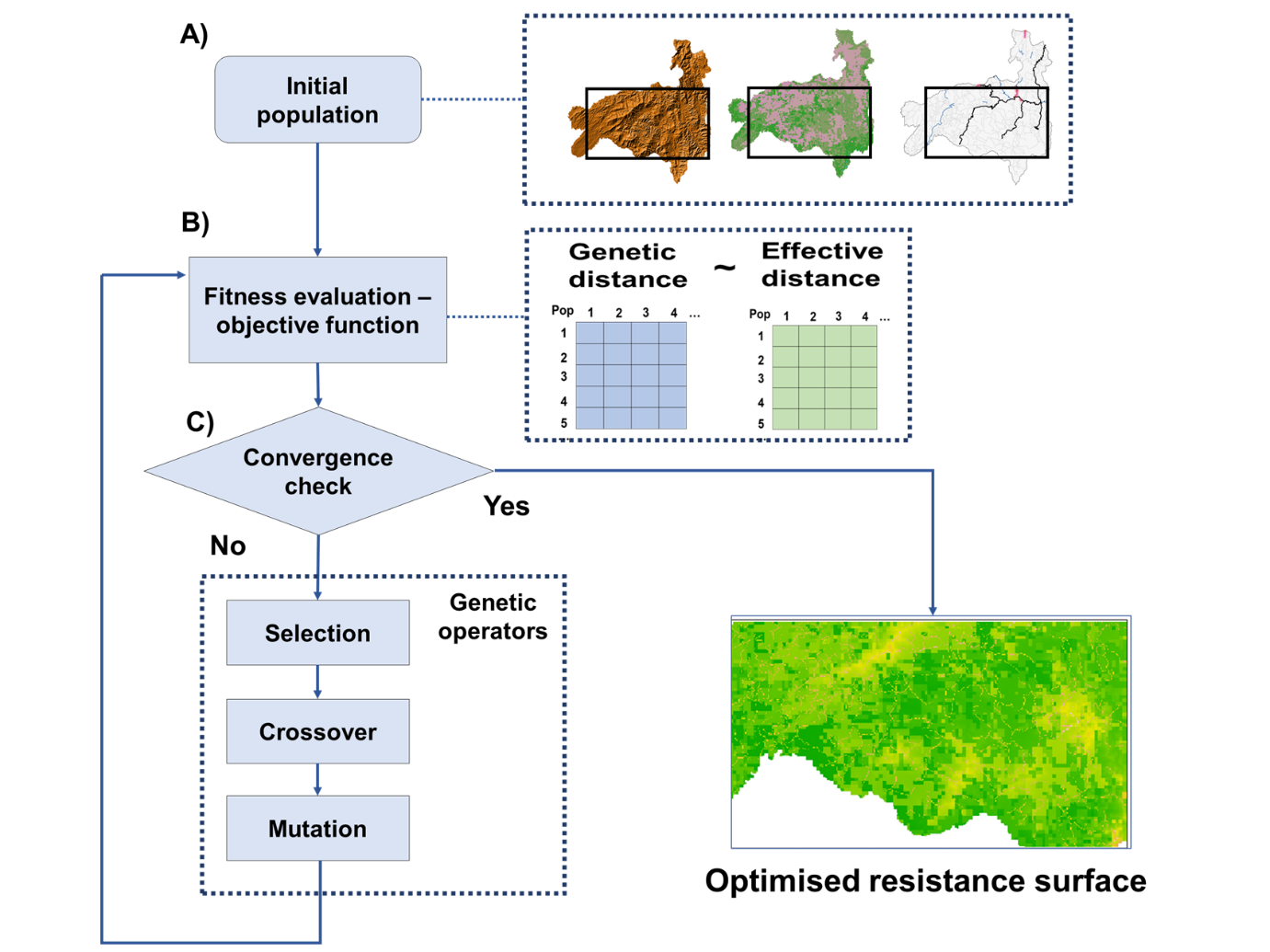

**Supplementary Figure 3 Schematic description of resistance surface optimisation in ResistanceGA. a,** The process starts with an initial population made of raster surfaces and their weighted parameters. **b,** An objective function is used to train in the GA algorithm. The objective function attempts to solve a linear mixed effects model with MLPE parametrisation that regress genetic and effective distances matrices. **c,** If the model fit cannot be improved, the process terminates, otherwise it continues finding the fittest individuals and weighted parameters using an evolvability process. This process selects (selection) the best individuals, “parents”, and reproduces (crossover) them to create a new generation with the best weighted parameters but slightly changed at random (mutation). The output of the ResistanceGA process is an optimised resistance surface with the optimal parameters solution.

In our analysis, effective distances (represented as commute distances) between sample sites locations (nodes) were calculated through random-walk commute time algorithm^25^ on our optimised surface (Supplementary Figure 4a) using the commuteDistance function implemented in the gdistance^26^ R package. Briefly, commute distance (Supplementary Figure 4b) can be defined as the expected average time a random walker travels, back-and-forth, between two nodes through a set of paths, which is analogous to an electrical circuit. Commute distance is an equivalent to resistance distance, but the latter accounts for passage cost and availability of alternative paths. Resistance distances can also be calculated in ResistanceGA by calling the circuit theory algorithm^27,28^ (Figure 4-8 C) of CIRCUITSCAPE v5^29^ software, however, computation is slightly slower than using commuteDistance.

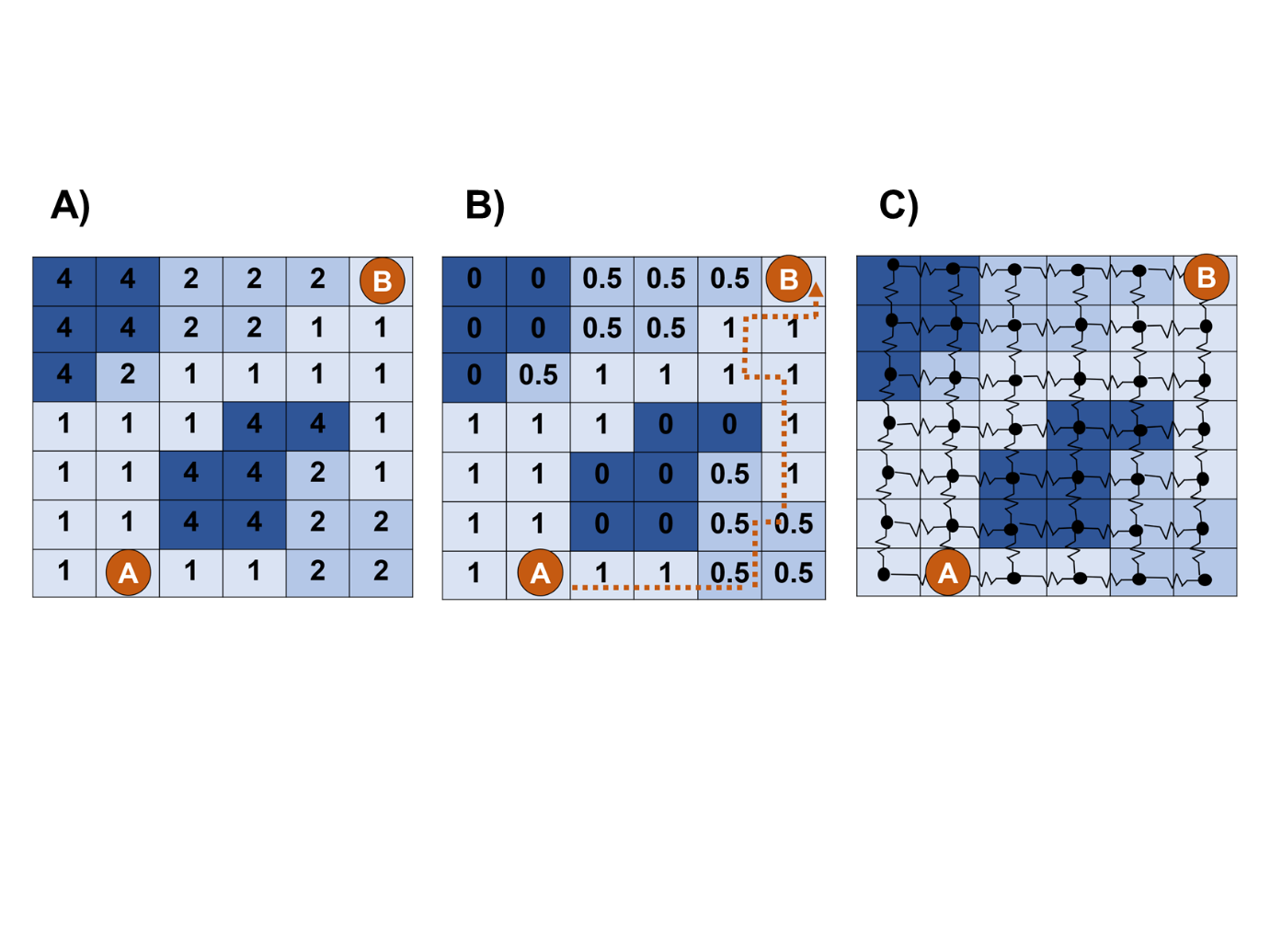

**Supplementary Figure 4 Schematic representation of random-walk commute time and circuit theory algorithms. a,** Raster surface with each grid cell provided with a resistance value (per-cell resistance increases with darker colour and values) based on the ResistanceGA optimisation process. Orange circles are focal nodes, labelled as A and B, representing sampling locations. **b**,	In the random-walk commute time algorithm, the raster grid is converted into a transition matrix with, *p*, probability that a random walker will step on that cell. Probabilities are inversely proportional to the resistance values in grid cells in **a**. Commute distance is calculated from the expected time this random walker will travel from focal node A to focal node B, back and forth, through a set of paths^26^. **c**, In circuit theory, raster grids are converted into a circuit network in which nodes (black dots) are connected by edges with resistors weighted inversely proportional to the resistance values in grid cells in **a**. If we apply a 1-amp current source to focal node A, current will flow through all nodes connected by weighted resistors until it reaches a grounded focal node B. Resistance distance is the accumulative passage of this current through these resistors averaged from the total available paths^27^.

**Supplementary Figures.**

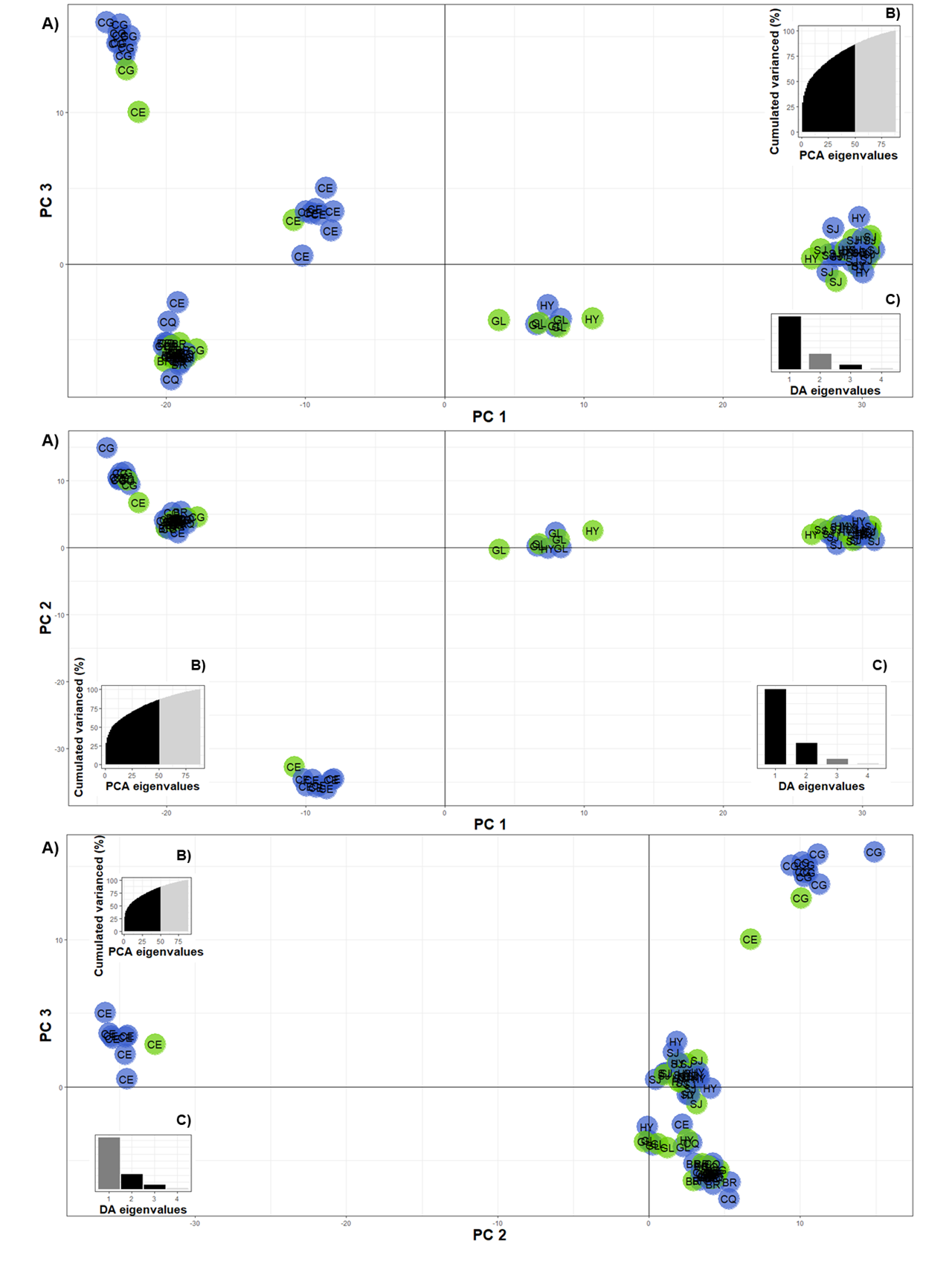

**Supplementary Figure 5 Discriminant analysis of principal components (DAPC) scatter plots of all possible PCs axes combinations comparing ecotope vs collection site in 89 samples using 2,552 SNP markers.** **a,** The scatter plots show different PCs combinations of the discriminant analysis eigenvalues for this small dataset. Five clusters can be seen in the scatter plots with dots representing individual samples coloured-coded to indicate their domestic (blue) or wild (green) ecotope of collection. Dots 4-digit label indicate the community of collection. Insets show **b** the percentage of cumulated variance explained (86%) by the PCs retained (black area) in the DAPC and **c** the different combinations of eigenvalues (black bars) for the discriminant functions of the discriminant analysis.

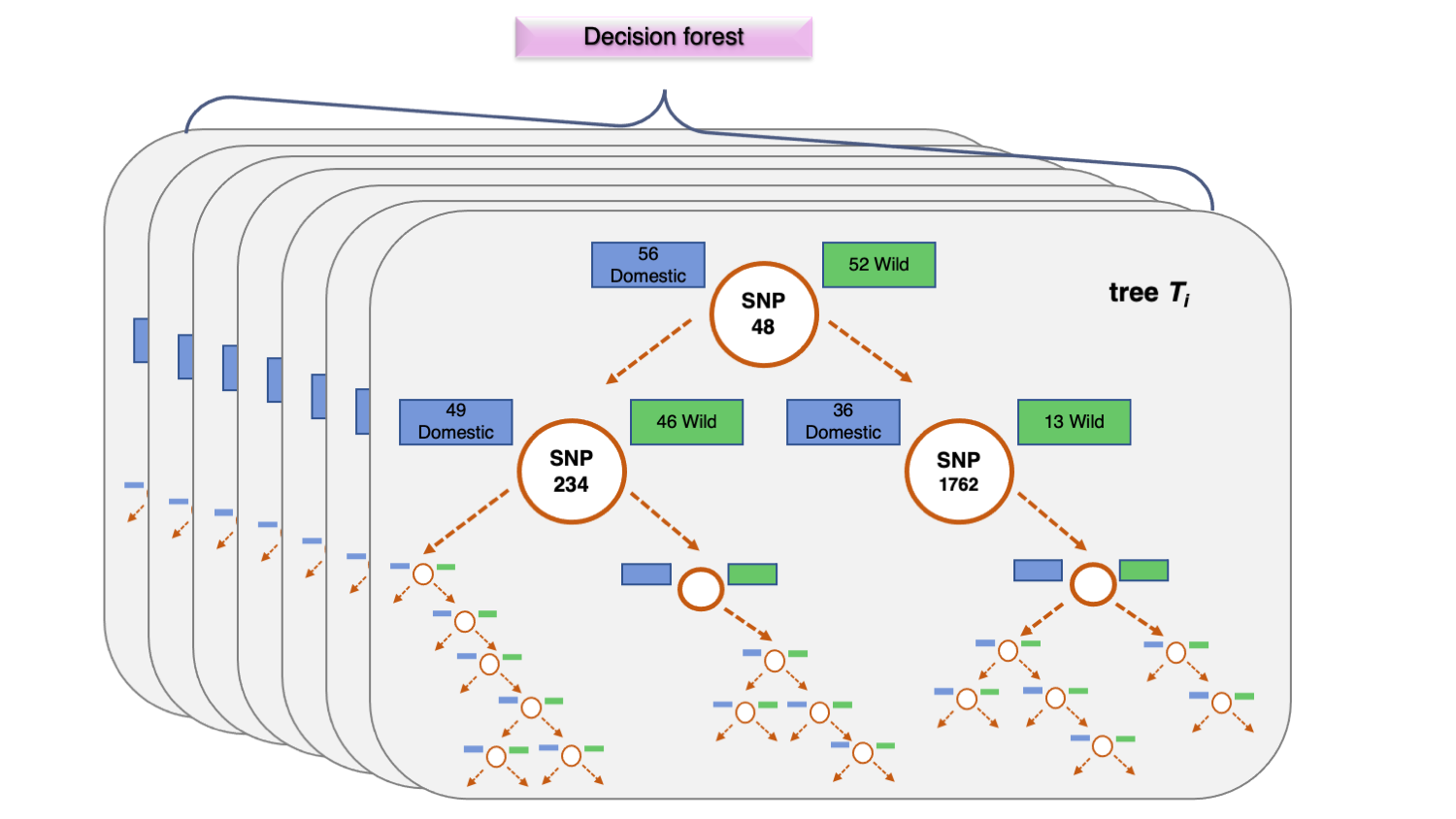

**Supplementary Figure 6 Random Forest procedure.** The algorithm starts training by building a n number of decision trees, a forest, with 66.6% of the samples. Each decision tree attempts to find the best SNPs that explain most of the within group variation (e.g. domestic and wild groups) until it is fully grown. Once trained, it compares the results with the remaining 33.3% of the samples and the misclassification of samples, or our-of-bag error rate, is used to find the SNPs with the best classification power. Boxes are colour-coded to represent domestic (blue) and wild (green) samples. SNPs and SNPs identification number are shown inside white circles. Each grey rectangle large box represents a decision tree within a large decision forest. This figure was adapted from https://dimensionless.in/introduction-to-random-forest/ and ^30^.

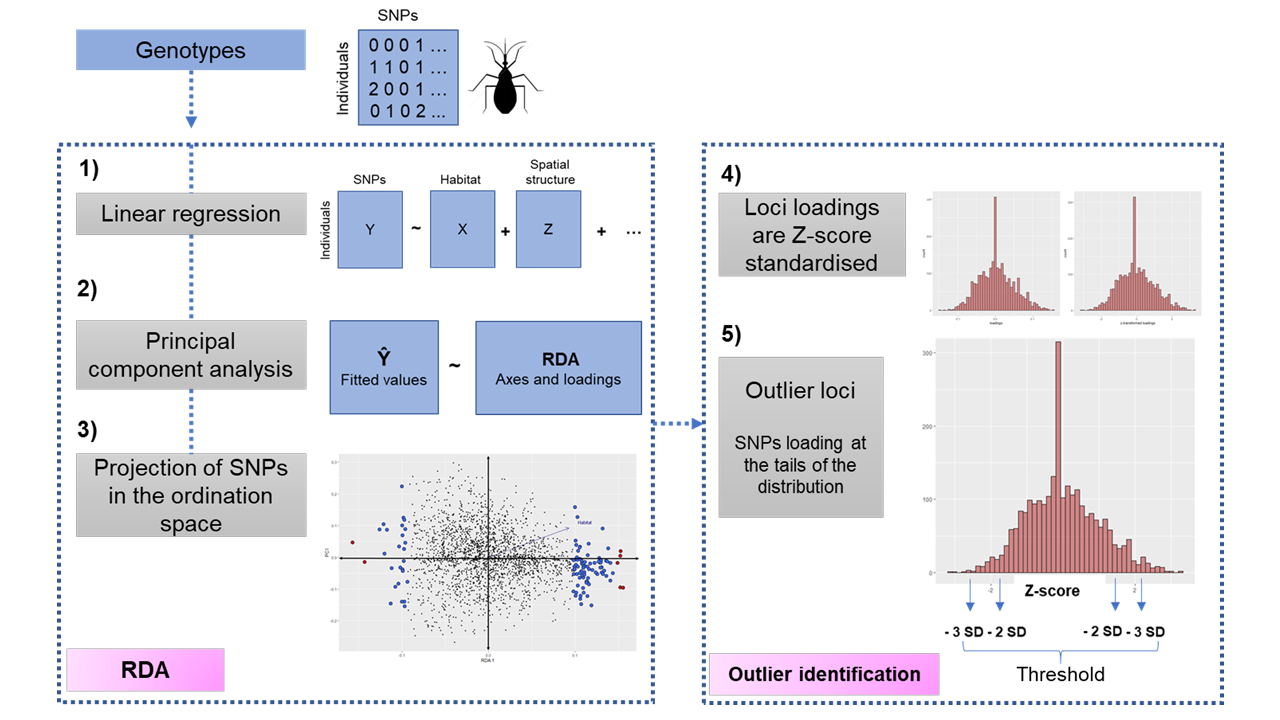

**Supplementary Figure 7 Step-by-step process of redundancy analysis (RDA).** The RDA performs, **a**, multivariate regression between a response matrix of genotypes, Y, and a matrix of explanatory variables, X. In our case, we added a conditional term to control for spatial structure, Z, based on the axes on individual principal coordinates of each sample. Then, a principal components analysis is carried out, **b**, on the matrix of fitted values, Ŷ, estimated from the regression on each locus. **c**, Relationship between SNPs coordinates and RDA axis can be explored in the ordination space. To identify outlier loci, **d**, SNPs loadings are Z- score standardised and, **e**, loci that fall at the tails of the distribution at a determine threshold (±2 SD or ±3 SD) are considered as outlier. This figure was adapted from ^31^.

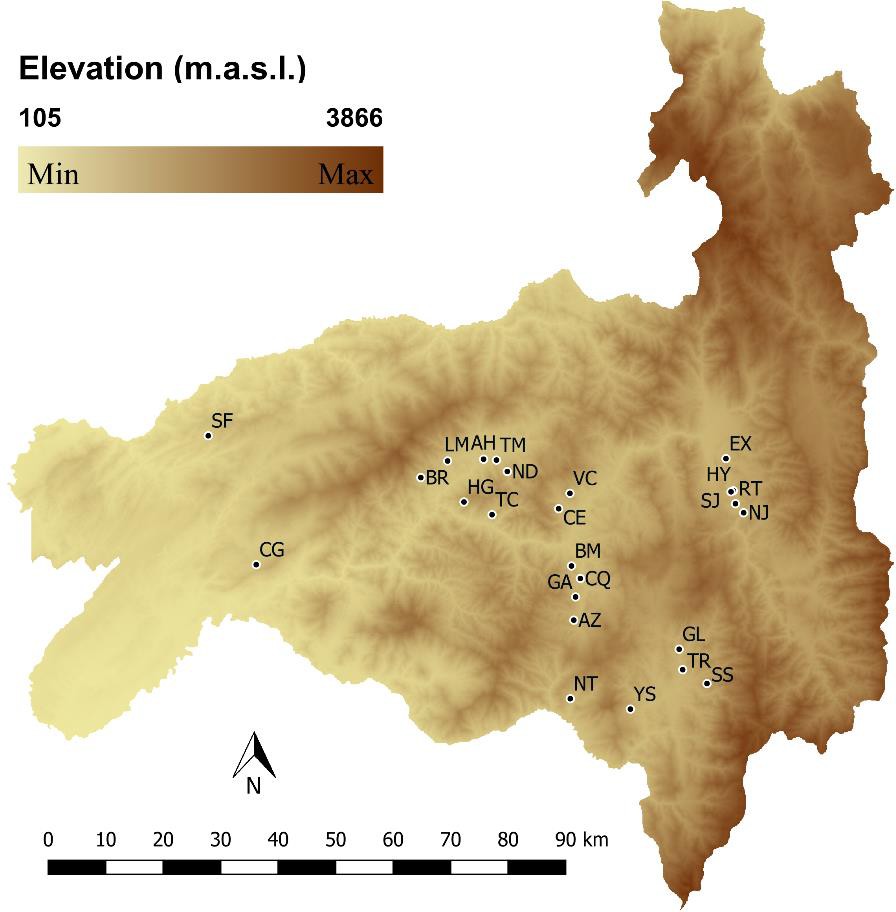

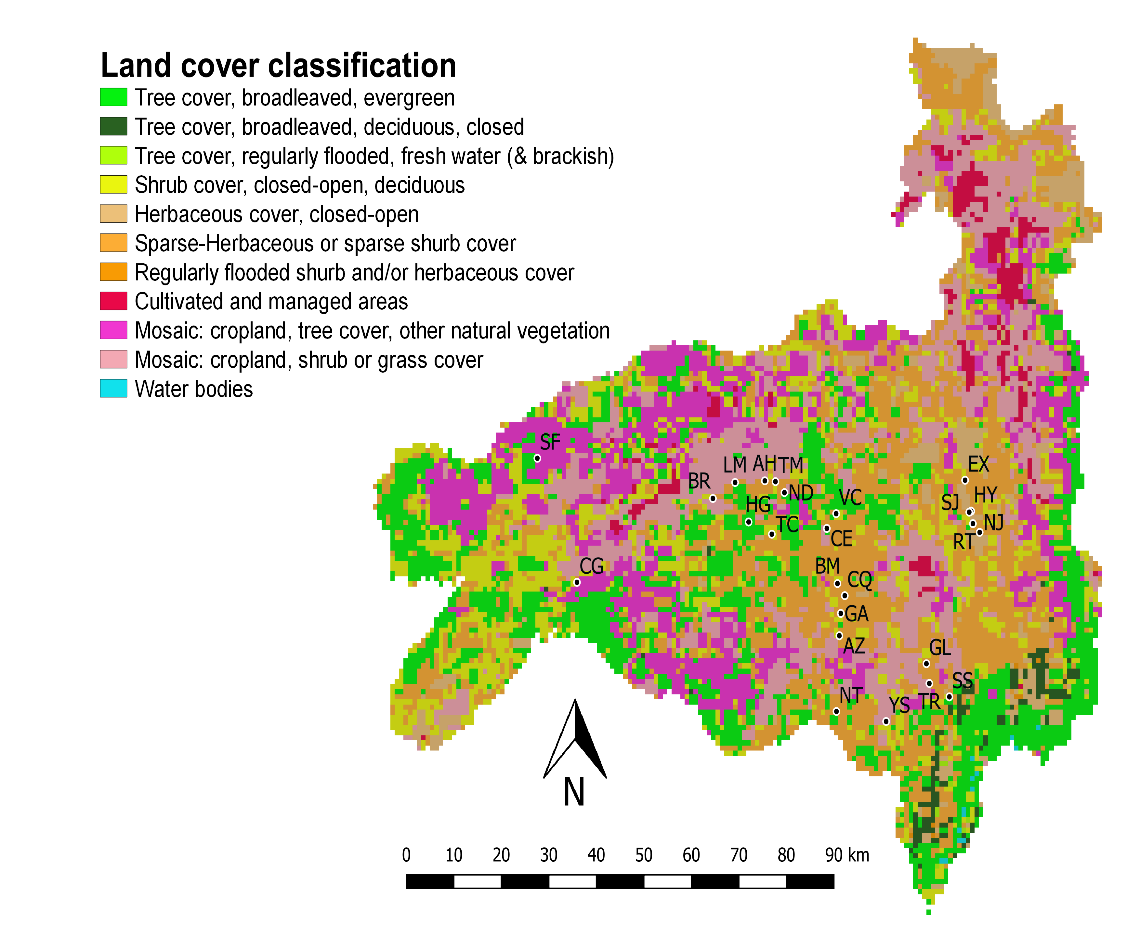
**Supplementary Figure 8 Relief of Loja, Ecuador.** Dots show the location of the 25 sampled communities at different altitudes across Loja. Relief is represented by a digital elevation model of brown colour gradient showing higher altitudes in dark shades of brown. Data from the SRTM database at <http://srtm.csi.cgiar.org/>.

**Supplementary Figure 9 Map of the land cover types present in Loja, Ecuador.** Dots show the 25 communities sampled in Loja at different land cover types (Supplementary Table 7). Different colours indicated one of the eleven single GLC2000 classification categories present in Loja. Data collected from the DIVA-GIS programme at https://[www.diva-gis.org/.](http://www.diva-gis.org/)

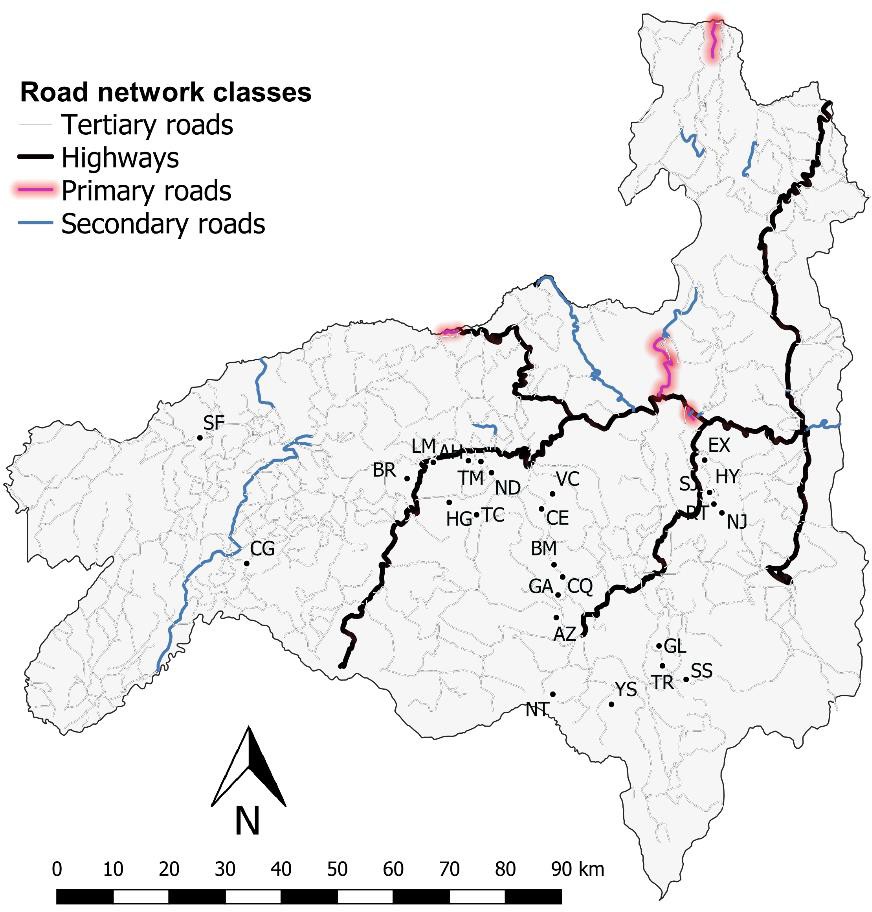

**Supplementary Figure 10 Map of the road network of Loja, Ecuador.** Dots show the location of the 25 sampled communities in Loja. Road categories are colour-coded to represent tertiary (grey thin line), highways (bold black line), primary (highlighted red) and secondary (light blue thin line) roads. Data collected from the GRIP dataset^32^ at www.globio.info/download-grip-dataset.

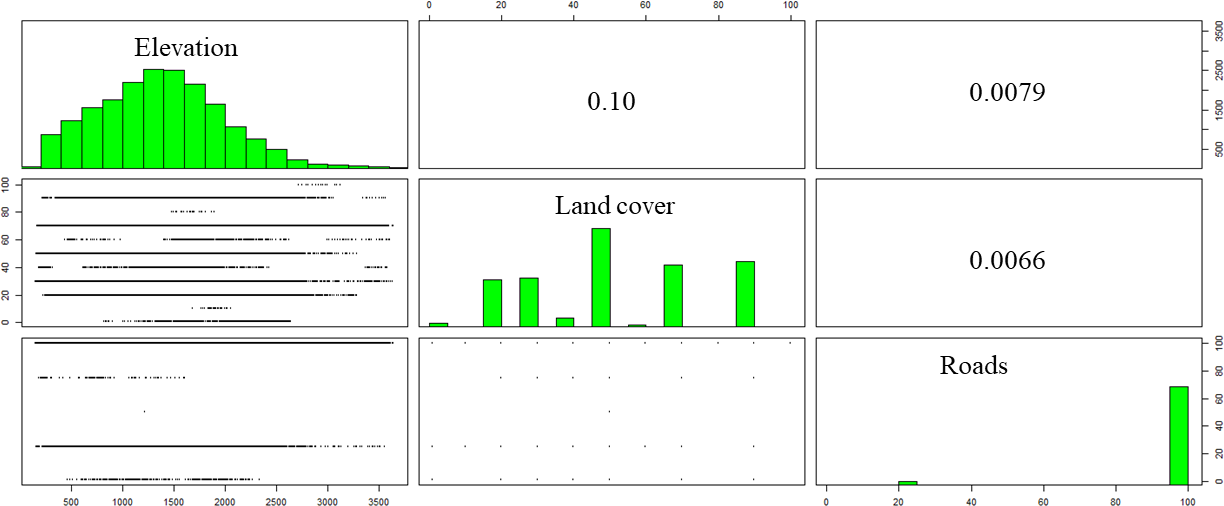

**Supplementary Figure 11 Scatterplot matrix showing the relation between raster surfaces.** Low correlation between relief-land cover (0.10), relief-roads (0.0079) and land cover-roads (0.0066) is observed.

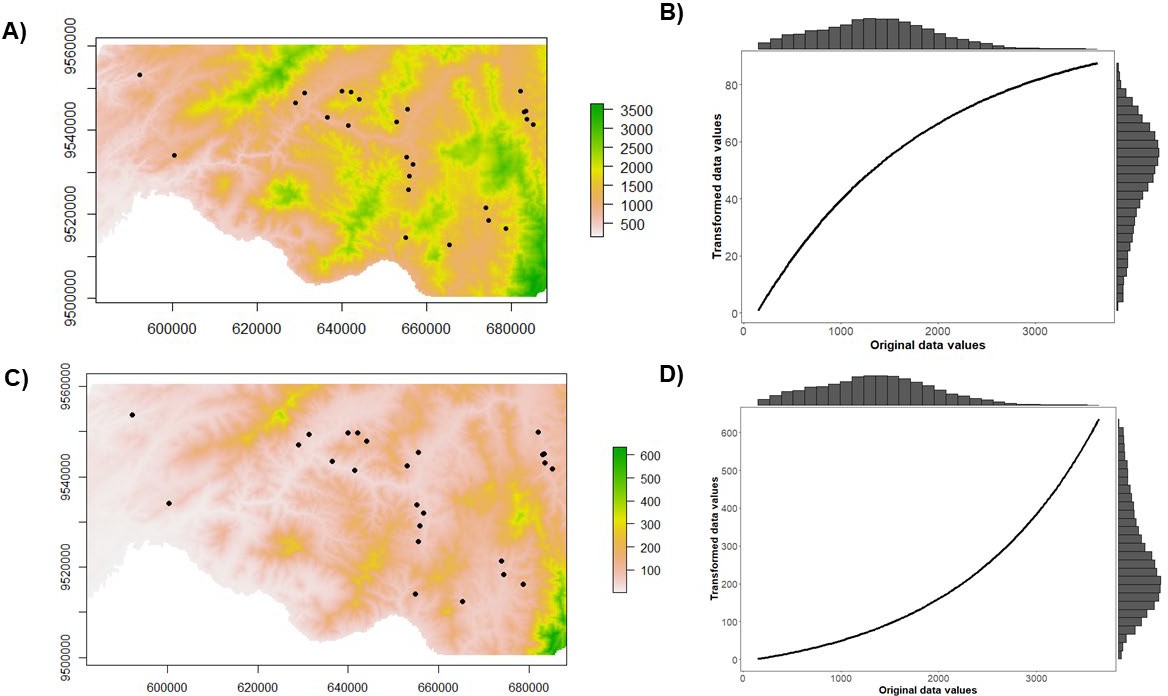

**Supplementary Figure 12 Comparison between original and optimised relief resistance surfaces**. Dots in **a** and **c** represent the sample location coordinates in UTM. Histograms in **b** and **d** represent the frequency of original and transformed altitude values. **a,** original altitude values of the study area. **b**, our initial hypothesis suggested an increase of cost of movement as altitude increase (monomolecular transformation) with the higher resistance values above 2,000 m.a.s.l. **c**, Resistance surface for relief showing optimised resistance values. **d**, Relief is suggested to follow a monotonical increase of cost of movement as relief increase (inverse-reverse transformation) with the highest resistance at approximately under 2,414 m.a.s.l.

**Supplementary Tables.**

**Supplementary Table 3 *Rhodnius ecuadoriensis* population genetic summary statistics.** Community name (Collection site), 2-letter ID labels (Code), sample collection ecotope (ecotope), number of samples (n), mean observed heterozygosity (HO), mean gene diversity (H_E_), inbreeding index (F_IS_) and allelic richness (A_r_) is provided. A chi-square test based on the expected genotype frequencies was used to calculated % HWE which is the proportion of loci with a p-value > 0.05 (* indicated the lowest proportions in HWE). Bold H_O_, H_E_, F_IS_ and A_r_ values inside yellow boxes indicated significant differences existed between domestic and wild mean diversity according to randomisation and t-test permutations.

| ***Collection site*** | ***Code*** | ***ecotope*** | ***n*** | ***H_O_*** | ***H_E_*** | ***FIS*** | ***A_r_*** |
| --- | --- | --- | --- | --- | --- | --- | --- |
| *San*  *Francisco* | SF | domestic | 6 | 0.21 | 0.17 | -0.19 | 1.35 |
| *La Cienega* | CG | domestic | 9 | **0.21** | **0.21** | **0.01** | 1.36 |
|  |  | wild | 4 | **0.23** | **0.22** | **-0.05** | 1.38 |
| *Bramaderos* | BR | domestic | 7 | **0.16** | **0.17** | **0.10** | **1.30** |
|  |  | wild | 8 | **0.23** | **0.22** | **-0.03** | **1.39** |
| *Limones* | LM | domestic | 11 | 0.20 | 0.21 | 0.04 | 1.44 |
| *Ashimingo* | AH | domestic | 18 | 0.20 | 0.19 | 0.01 | 1.40 |
| *Naranjo Dulce* | ND | domestic | 15 | 0.19 | 0.17 | -0.04 | 1.37 |
| *Higida* | HG | domestic | 10 | 0.17 | 0.16 | -0.02 | 1.35 |
| *Tacoranga* | TC | domestic | 10 | 0.16 | 0.18 | 0.11 | 1.37 |
| *Vega del Carmen* | VC | domestic | 11 | 0.15 | 0.16 | 0.10 | 1.34 |
| *Coamine* | CE | domestic | 9 | **0.16** | **0.16** | **0.05** | **1.28** |
|  |  | wild | 2 | **0.19** | **0.21** | **-0.06** | **1.31** |
| *Bella Maria* | BM | domestic | 3 | 0.22 | 0.21 | -0.08 | 1.41 |
| *Chaquizhca* | CQ | domestic | 9 | 0.18 | 0.19 | **0.08** | **1.34** |
|  |  | wild | 2 | 0.17 | 0.18 | **-0.09** | **1.27** |
| *Guara* | GA | domestic | 10 | 0.22 | 0.19 | -0.08 | 1.41 |
| *La Extensa* | EX | domestic | 20 | 0.11 | 0.09 | -0.09 | 1.19 |
| *San Jacinto* | SJ | domestic | 10 | 0.14  0.14 | 0.12  0.12 | -0.12  -0.13 | 1.21  1.21 |
|  |  | wild 9 | |  |  |  |  |
| *El Huayco* | HY | domestic | 10 | **0.12**  **0.17** | **0.12**  **0.15** | **-0.02**  **-0.17** | **1.21**  **1.25** |
|  |  | wild 3 | |  |  |  |  |
| *Camayos* | YS | domestic | 19 | 0.19 | 0.16 | -0.08 | 1.34 |
| *Galapagos* | GL | domestic | 3 | **0.16**  **0.20** | **0.16**  **0.21** | **-0.08**  **0.022** | **1.26**  **1.35** |
|  |  | wild 4 | |  |  |  |  |
| *Tuburo* | TR | domestic | 8 | 0.17 | 0.16 | -0.03 | 1.34 |
| *Santa Rosa* | SS | domestic | 10 | 0.14 | 0.15 | 0.09 | 1.32 |
| *Naranjillo* | NJ | domestic | 4 | 0.19 | 0.19 | -0.03 | 1.39 |
| *Ardanza* | AZ | wild | 8 | 0.17 | 0.18 | 0.03 | 1.37 |
| *San Antonio de Taparuca* | NT | wild | 9 | 0.15 | 0.12 | -0.24 | 1.25 |
| *Santa Rita* | RT | wild | 6 | 0.13 | 0.11 | -0.13 | 1.23 |
| *Tamarindo* | TM | domestic | 5 | 0.15 | 0.14 | -0.03 | 1.29 |

**Supplementary Table 4 Summarised results of hierarchical differentiation of molecular variance components in 2552 SNP loci on the complete (n = 272 samples) and a small (n = 89 samples) datasets.** Variance components, F-statistic (= G-statistic^33^), confidence interval (C.I. at 2.5% and 97.5%) and P-values (significance level based in randomised 999 permutation of individuals level-defined) are provided.

| ***Source of*** | |  | **dataset (n = 272)** | |  | **dataset (n = 89)** | |  |  |
| --- | --- | --- | --- | --- | --- | --- | --- | --- | --- |
| ***variation*** | | Variance component | F-statistic | C.I. | p-value | Variance F-statistic components | | C.I, | p-value |
| *Among communities* | | 151.21 | 0.26 | 0.25  0.27 | 0.001 | 149.91 | 0.25 | 0.24  0.26 | 0.001 |
| *Among populations*  *within communities* | | -18.70 | -0.04 | -0.05  -0.04 | 0.001 | -2.04 | -0.004 | -0.01  0.004 | 0.001 |
| *Among collection year within*  *communities* | | 52.19 | 0.07 | 0.07  0.08 | 0.001 | 30.06 | 0.06 | 0.058  0.067 | 0.001 |
| *Individuals*  *communities* | *within* | -34.19 | -0.001 | -0.02  0.02 |  | -18.13 | 0.02 | -0.001  0.04 |  |
| *Error* | | 436.63 | | | | 436.01 | | | |

**Supplementary Table 5 GLS-MLPE model results.** Estimated regression parameters with 95% confidence intervals (Lower/Upper), standard errors, t-values and p-values for the GLS- MLPE model presented in eqn 1.

|  | **Estimate** | **Lower** | **Upper** | **Std. error** | **t value** | **P‐value** |
| --- | --- | --- | --- | --- | --- | --- |
| **Intercept** | 0.12 | 0.08 | 0.16 | 0.021 | 5.67 | < 0.001 |
| **Geographic**  **distance** | 0.0016 | 0.0014 | 0.0018 | 0.0001 | 14.13 | < 0.001 |
| **Ecotope Wild** | -0.07 | -0.15 | 0.06 | 0.04 | -1.88 | 0.06 |
| **Geographic distance: Ecotope Wild** | 0.0018 | 0.0011 | 0.0024 | 0.003 | 5.72 | < 0.001 |

**Supplementary Table 6 Original GIS data, description and collection source.**

| **Original GIS data** | **Original format** | **Source** |
| --- | --- | --- |
| Relief was obtained from a digital elevation model | - Raster format. - Resolution of 90 m at the equator. - Minimum altitude of 105 and maximum of 3866 m.a.s.l. | Shuttle Radar Topography Mission (SRTM) database (<http://srtm.csi.cgiar.org/>) |
| Land cover | - Raster format - Resolution approximately 1 Km. - Eleven distinct land cover classifications. - Classification is based on the Global Land Cover (GLC) 2000 project which follows United Nations Food and Agricultural Organisation (FAO) Land Cover Classification System (LCCS). Supplementary Table 7 | DIVA-GIS programme (https://www.diva-gis.org/) |
| Road network | - Vector format. - Five road categories: highways, primary, secondary, tertiary and no roads. | Global Roads Inventory Project (GRIP) dataset^32^ (www.globio.info/ download-grip-dataset). |

**Supplementary Table 7 Global Land Cover 2000 project Legend.** Aggregated from regional classes using United Nations Food and Agricultural Organisation (FAO) Land Cover Classification System (LCCS). DIVA-GIS programme at https://www.diva-gis.org/.

|  | **GLC Global Class (according to LCCS terminology)** |
| --- | --- |
| 1 | **Tree Cover, broadleaved, evergreen**  *LCCS >15% tree cover, tree height >3m*  (Examples of sub-classes at regional level* :  *closed > 40% tree cove; open 15-40% tree cover)* |
| 2 | **Tree Cover, broadleaved, deciduous, closed** |
| 3 | **Tree Cover, broadleaved, deciduous, open**  *(open 15-40% tree cover)* |
| 4 | **Tree Cover, needle-leaved, evergreen** |
| 5 | **Tree Cover, needle-leaved, deciduous** |
| 6 | **Tree Cover, mixed leaf type** |
| 7 | **Tree Cover, regularly flooded, fresh water** (& brackish) |
| 8 | **Tree Cover, regularly flooded, saline water,**  **(**daily variation of water level) |
| 9 | **Mosaic:**  **Tree cover / Other natural vegetation** |
| 10 | **Tree Cover, burnt** |
| 11 | **Shrub Cover, closed-open, evergreen**  (Examples of sub-classes at reg. level *: (i) sparse tree layer) |
| 12 | **Shrub Cover, closed-open, deciduous**  (Examples of sub-classes at reg. level *: (i) sparse tree layer) |
| 13 | **Herbaceous Cover, closed-open**  (Examples of sub-classes at regional level *:  (i) natural, (ii) pasture, (iii) sparse trees or shrubs) |
| 14 | **Sparse Herbaceous or sparse Shrub Cover** |
| 15 | **Regularly flooded Shrub and/or Herbaceous Cover** |
| 16 | **Cultivated and managed areas**  (Examples of sub-classes at reg. level *:  (i) terrestrial; (ii) aquatic (=flooded during cultivation), and under terrestrial: (iii) tree crop & shrubs (perennial), (iv) herbaceous crops (annual), non-irrigated, (v) herbaceous crops (annual), irrigated) |
| 17 | **Mosaic:**  **Cropland / Tree Cover / Other natural vegetation** |
| 18 | **Mosaic:**  **Cropland / Shrub or Grass Cover** |
| 19 | **Bare Areas** |
| 20 | **Water Bodies** (natural & artificial) |
| 21 | **Snow and Ice** (natural & artificial) |
| 22 | **Artificial surfaces and associated areas** |

**Supplementary Table 8 Spearman correlation test, rho, between raster surfaces previous to optimization with ResistanceGA.**

| **Surface 1** | **Surface 2** | **Spearman’s rho** |
| --- | --- | --- |
| Relief | Land cover | -0.1315174 |
| Relief | Roads | 0.01405202 |
| Roads | Land cover | 0.03044303 |

**Supplementary Table 9 Land cover reclassified values.** Original and reclassified resistance values for the different land cover categories are provided.

| **GLC GLOBAL CLASS (ACCORDING TO**  **LCCS TERMINOLOGY)** | **ORIGINAL**  **VALUE** | **TRANSFORMED**  **VALUE** |
| --- | --- | --- |
| **TREE COVER, BROADLEAVED, EVERGREEN** | 1 | 70 |
| **TREE COVER, BROADLEAVED, DECIDUOUS, CLOSED** | 2 | 60 |
| **TREE COVER, REGULARLY FLOODED, FRESH WATER (& BRACKISH)** | 7 | 90 |
| **SHRUB COVER, CLOSED-OPEN, DECIDUOUS** | 12 | 30 |
| **HERBACEOUS COVER, CLOSED-OPEN** | 13 | 40 |
| **SPARSE HERBACEOUS OR SPARSE SHRUB**  **COVER** | 14 | 50 |
| **REGULARLY FLOODED SHRUB AND/OR**  **HERBACEOUS COVER** | 15 | 80 |
| **CULTIVATED AND MANAGED AREAS** | 16 | 1 |
| **MOSAIC: CROPLAND / TREE COVER / OTHER NATURAL VEGETATION** | 17 | 20 |
| **MOSAIC: CROPLAND / SHRUB OR GRASS COVER** | 18 | 10 |
| **WATER BODIES (NATURAL & ARTIFICIAL)** | 20 | 100 |

**Supplementary Table 10 Roads reclassified values.** Original and reclassified resistance values for the different road classes are provided.

| **TERTIARY ROADS** | 0 | 25 |
| --- | --- | --- |
| **HIGHWAYS** | 1 | 1 |
| **PRIMARY**  **ROADS** | 2 | 50 |
| **SECONDARY ROADS** | 3 | 75 |
| **NO ROADS** | 4 | 100 |

**Supplementary References.**
